# Activity-dependent netrin-1 secretion drives synaptic insertion of GluA1-containing AMPA receptors in the hippocampus

**DOI:** 10.1101/330688

**Authors:** Stephen D. Glasgow, Simon Labrecque, Ian V. Beamish, Sarah Aufmkolk, Julien Gibon, Dong Han, Stephanie N. Harris, Paul Dufresne, Paul W. Wiseman, R. Anne McKinney, Philippe Séguéla, Paul De Koninck, Edward S. Ruthazer, Timothy E. Kennedy

## Abstract

Dynamic trafficking of α-amino-3-hydroxy-5-methyl-4-isoxazolepropionic acid glutamate receptors (AMPARs) to synapses is critical for activity-dependent synaptic plasticity underlying learning and memory, however the identity of key molecular effectors remains elusive. Here, we demonstrate that membrane depolarization and N-methyl-D-aspartate receptor (NMDAR) activation triggers secretion of the chemotropic guidance cue netrin-1 from dendrites. Using selective genetic deletion, we show that netrin-1 expression by excitatory neurons is required for NMDAR-dependent long-term potentiation (LTP) in the adult hippocampus. Further, we demonstrate that application of exogenous netrin-1 is sufficient to trigger the potentiation of excitatory glutamatergic transmission at hippocampal Schaffer collateral synapses via Ca^2+^-dependent recruitment of GluA1-containing AMPARs, promoting the maturation of immature or nascent synapses. These findings identify a central role for activity-dependent release of netrin-1 as a critical effector of synaptic plasticity in the adult hippocampus.

---

Glutamatergic synaptic transmission in the adult brain is primarily mediated by AMPARs, with changes in synaptic strength due to alterations in both synaptic structure and receptor composition ^1, 2^. Long-term potentiation (LTP), an intensively studied experimental model of synaptic plasticity and memory, modifies postsynaptic function through NMDAR-dependent recruitment of AMPARs, cytoskeletal reorganization, and modification of synaptic adhesion ^3-6^.

Netrin-1 is a secreted protein that directs axon outgrowth and synaptogenesis during development ^7-9^. The netrin receptor, deleted in colorectal cancer (DCC), directs cell motility by regulating intracellular calcium, RhoGTPases, Src family kinases, focal adhesion kinase, phospholipase C, phosphoinositol 3-kinase, p21-activated kinase, and local protein synthesis, all of which influence adhesion, cytoskeletal organization, and synapse function ^10, 11^. During neural development, restricted sources of netrin-1 direct the rapid local recruitment of synaptic proteins ^7, 12, 13^. Further, a form of synaptic consolidation in *Aplysia* neurons, studied in primary cell culture, requires netrin-1-dependent local protein synthesis, suggesting a critical role for netrin-1 in the modification of synaptic function ^14^. In the mature mammalian brain, netrin-1 and DCC are enriched at synapses, and DCC co-fractionates with detergent resistant components of the postsynaptic density (PSD). Selective deletion of DCC from excitatory forebrain neurons in the mature brain reduces dendritic spine volume, severely attenuates LTP at Schaffer collateral hippocampal synapses, and impairs hippocampal-dependent learning and memory ^15^. Several families of ligands have been identified for DCC, including draxins, cerebellins, and netrins ^16-18^; however, the enrichment of netrin-1 at synapses suggests that it may be a key ligand for DCC to regulate synaptic function.

Here, we show that activity-dependent secretion of netrin-1 from neurons regulates synaptic transmission in the adult hippocampus. Selective genetic deletion of netrin-1 from principal excitatory neurons severely attenuates NMDAR-dependent LTP in the adult mammalian hippocampus. Further, application of exogenous netrin-1 to Schaffer collateral synapses is sufficient to induce a lasting potentiation of evoked synaptic responses through the recruitment of GluA1-containing, GluA2-lacking AMPARs. Taken together, our study identifies the activity-dependent secretion of netrin-1 as a critical component of LTP expression downstream of NMDAR activation.

## Results

### Netrin-1 is secreted in response to neuronal depolarization and NMDAR activation

Netrin-1 is highly enriched in intracellular vesicles in synaptosomes ^15^, suggesting that action potential generation and neuronal depolarization might regulate netrin-1 release at synapses. To initially examine if netrin-1 protein may be secreted from neurons in an activity-dependent manner, we collected and concentrated cell-culture supernatants of embryonic rat neurons (14 DIV) that had been depolarized for 20 min with 20 mM KCl. Immunoblots of the concentrated supernatants revealed significantly increased levels of netrin-1 compared to supernatants from cultures treated with vehicle alone (Supplemental figure 1), indicating that netrin-1 is secreted in response to strong depolarization.

Sustained neuronal activity can mobilize vesicular release ^19^. To determine if increased neuronal spiking induces secretion of netrin-1, we used the designer receptor exclusively activated by designer drugs (DREADD) approach to excite primary hippocampal neurons infected with AAV8-Syn-hM3D(Gq)-mCitrine. Bath application of the synthetic hM3D(Gq) agonist clozapine-N-oxide (CNO; 10 µM) triggered rapid depolarization and spiking of mCitrine-expressing neurons (Supplemental figure 1). We detected a large increase in netrin-1 protein in supernatants from cultures expressing hM3D compared to untransfected control cultures following bath application of CNO (10 µM) for 1 h (Figure 1a), demonstrating that prolonged spiking and depolarization of hippocampal neurons promotes secretion of netrin-1.

**Figure 1.**
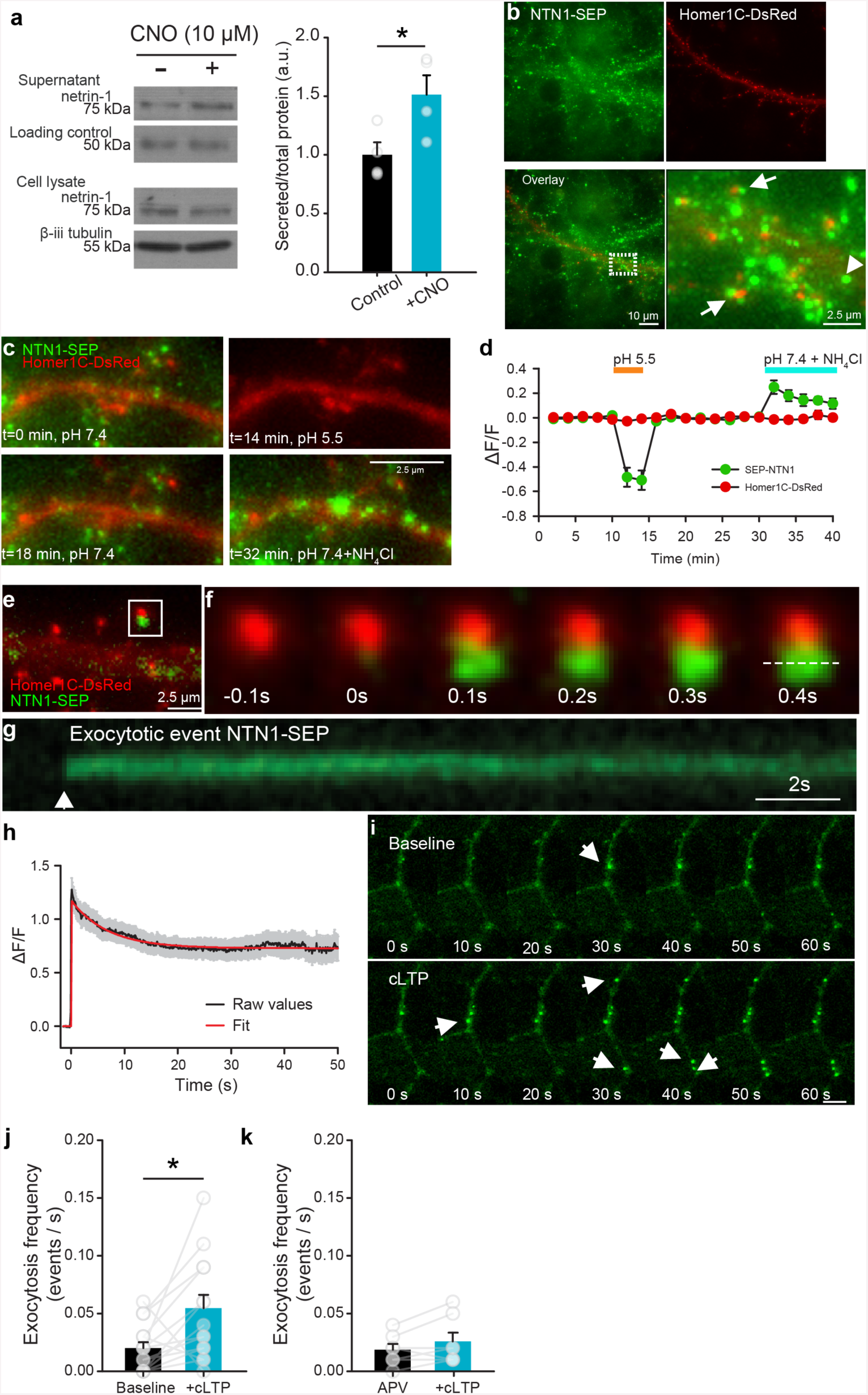
Netrin-1 is secreted by neurons in response to depolarization and NMDAR activation. Western blots (left) and group data (right) of concentrated media supernatants and cell lysates of cultured hippocampal neurons (14 DIV) expressing hM3D(Gq) and treated with vehicle (black) or depolarized with CNO (10 µM, yellow) for 1h (control supernatant: 1.00±0.11, CNO supernatant: 1.51±0.17, *n* = 4, *t*_6_= 2.60, *p* = 0.04, independent samples *t*-test, *n* = 4 independent cultures per condition). Note that an Alexa 633-conjugated secondary antibody (40 ng per tube) was added to the medium as a loading control. (**b**) Low-(left panels) and high-magnification (far right) of NTN1-SEP (green) appears as puncta at synaptic sites (Homer1C-DsRed-positive, red) in cultured hippocampal neurons (14 DIV). (**c**) Representative segment of a hippocampal dendrite expressing NTN1-SEP (green) and Homer1C-DsRed (red) during baseline (0 min), following exposure to extracellular solution at pH 5.5 (14 min), recovery (18 min), and bath application of NH_4_Cl (32 min). (**d**) Group data show decrease in SEP-NTN1 fluorescence intensity following bath application of extracellular solution at pH 5.5 (orange line), and an increase at intra-dendritic sites following bath application of NH_4_Cl (blue line, *n* = 7 cells). (**e**) High magnification of Homer1C-DsRed (red) positive puncta showing NTN1-SEP (green). (**f)** Time-lapse of a Homer1C-DsRed puncta (red, in **e**) prior to and following a NTN1-SEP exocytotic event. (**g**) Kymograph of NTN1-SEP fluorescence of a Homer1C-DsRed puncta (dashed line in **f**). (**h**) Raw group data (average in black, SEM in grey) and fitted curve (red) show amplitude and duration of NTN1-SEP exocytotic events. (**i**) Time-lapse images of dendritic segment shown in **e** during baseline (top) and following cLTP induction (bottom). Note the NTN1-SEP exocytotic events (white arrows). (**j**) Group data show frequency of exocytotic NTN1-SEP events in during base-line and following cLTP induction (baseline: 0.020±0.005, cLTP: 0.055±0.011, *t*_13_=2.56, *p*=0.02, paired-samples *t*-test, *n*=14 dendritic segments from 14 independent cultures). (**k**) Group data show frequency of NTN1-SEP exocytosis during baseline and following cLTP induction in the presence of NMDAR antagonist APV (50 µM) (APV: 0.019±0.005, cLTP: 0.025±0.007, *t*_6_=1.70, *p*=0.14, *n*=7 dendritic segments from 7 independent cultures, paired-samples *t*-test).

The neuronal specific promoter (synapsin) used in the DREADD experiments indicates that depolarization causes netrin-1 to be released by neurons. However, whether netrin-1 is released perisynaptically by axons or dendrites remained unresolved. To reveal the subcellular localization of netrin-1 secretion, we developed a full-length functional netrin-1 fused to super-ecliptic pHluorin (NTN1-SEP) that increases fluorescence intensity upon shifting from acidic to neutral pH following vesicle fusion with the plasma membrane (Supplemental figure 2). Co-expression of NTN1-SEP and the synaptic marker Homer1C-DsRed in cultured rat hippocampal neurons (14 DIV) revealed NTN1-SEP positive puncta along dendrites at synaptic (Homer 1C-DsRed positive) and extrasynaptic sites (Figure 1b), in a distribution similar to endogenous netrin-1 (Supplemental figure 2). Fluorescence was quenched by a brief pulse of acidic solution (pH=5.5), confirming that it corresponds to extracellular secreted NTN1-SEP. Netrin-1 secreted from dendrites should be initially present in an intra-dendritic endosomal pool. Intracellular distributions of NTN1-SEP were detected by applying an extracellular NH_4_Cl solution, alkalinizing the intracellular compartment and revealing large clusters of NTN1-SEP throughout dendritic shafts and spines (Figure 1c-d).

To address whether netrin-1 secretion is constitutive or mobilized in response to synaptic activity, we performed wide-field fluorescence video imaging of NTN1-SEP in neurons co-transfected with Homer1C-DsRed. To detect exocytotic events, pre-existing cell-surface fluorescence was first photobleached ^20^. We then monitored the spontaneous appearance of new bright puncta that reached full fluorescence amplitude, within 1-2 imaging time points, consistent with rapid exocytosis of NTN1-SEP (Figure 1e-f). We observed that NTN1-SEP fluorescence decay frequently lasted for minutes, consistent with NTN1-SEP remaining extracellularly at synapses following secretion (Figure 1g-h).

NMDAR activation is critical for several major forms of activity-dependent synaptic plasticity ^21^. To determine if NMDAR activation drives netrin-1 secretion, we monitored the dynamic appearance of NTN1-SEP fluorescent puncta in hippocampal neurons following chemical LTP (cLTP), induced by applying an extracellular solution of 0 mM Mg^2+^, 200 µM glycine, and 30 µM bicuculline that promotes NMDAR conductance and potentiates AMPAR-mediated currents ^22, 23^. Inducing cLTP increased the frequency of NTN1-SEP exocytosis, and promoted rapid secretion of long-lasting NTN1-SEP puncta (Figure 1i-j, Supplemental movie 1). To examine whether the increase in frequency of NTN1-SEP exocytosis requires NMDAR activation, we repeated these experiments in the presence of 2-amino-5-phosphonovaleric acid (APV, 50 µM), a potent and specific NMDAR antagonist. cLTP stimulation in the presence of APV failed to increase the frequency of NTN1-SEP exocytotic events (Figure 1k), indicating that the increase in secretion requires NMDAR activation. Together, these findings demonstrate that NMDAR activation promotes netrin-1 secretion from dendrites.

### CA1 neurons in the adult hippocampus express netrin-1

RNA-seq analysis previously detected netrin-1 mRNA in adult mouse hippocampal CA1 neurons ^24^. Using a validated netrin-1 specific monoclonal antibody ^25^, immunohistochemical analyses of hippocampal sections from Thy1-GFP L15 mice revealed a punctate distribution of endogenous netrin-1 protein across all layers of the CA1 region, including strata oriens, pyramidale, and radiatum (Supplemental figure 3). These findings indicate that *ntn1* is expressed by hippocampal neurons, and that endogenous netrin-1 protein can be detected in all layers of the adult hippocampus.

The subcellular distribution of netrin-1 was then assessed throughout stratum radiatum in adult Thy1-GFP L15 mice using super-resolution confocal microscopy (Figure 2a, Supplemental figure 3). The apical dendrites of CA1 pyramidal neurons, visualized by membrane-targeted GFP expression, show punctate endogenous netrin-1 immunoreactivity throughout the neuropil (Supplemental figure 3). Immunostaining for netrin-1 shows clusters of endogenous protein throughout the dendritic shafts and spines. Many of these netrin-1 clusters are contained within dendritic spines, as revealed by opaque surface renderings of the membrane-targeted GFP (Figure 2a). Consistent with NTN1-SEP live imaging (Figure 1), these results localize endogenous netrin-1 protein within dendrites and spine heads of CA1 pyramidal neurons.

**Figure 2.**
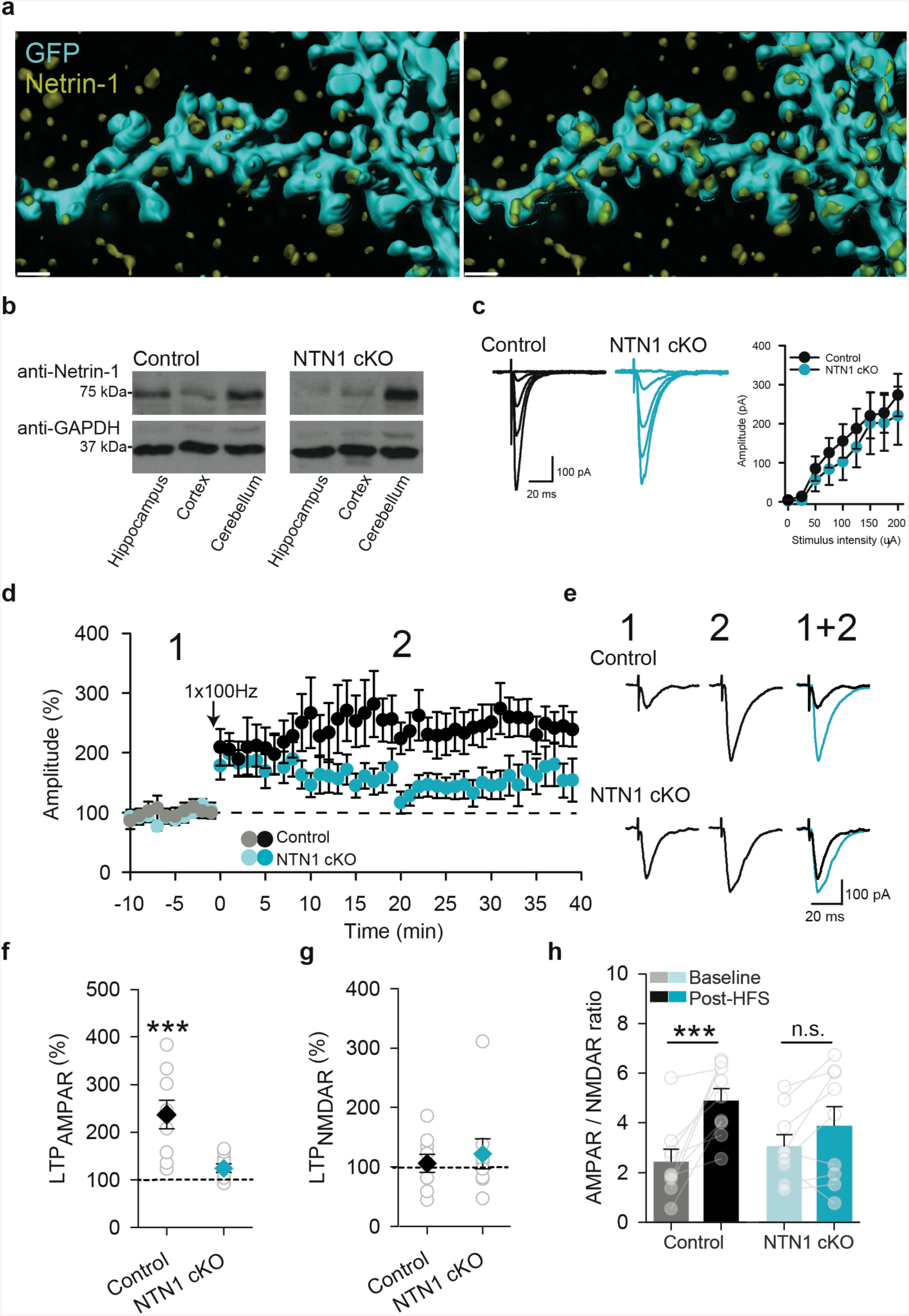
Neuronal netrin-1 is required for hippocampal LTP. (**a**) Dendrite of hippocampal CA1 neuron expressing GFP (cyan), counterstained with anti-netrin-1 (yellow). Full opacity (left) of membrane linked GFP shows netrin-1 immunoreactivity associated with the dendritic arbor. GFP transparency (right) reveals endogenous netrin-1 within the dendritic shaft and spines. Scale bar = 1 µm. (**b**) Representative western blots of whole hippocampal, cortical, and cerebellar homogenates from control wildtype littermates (left) and CaMKIIα∷Cre/NTN1^*fl/fl*^ (right) mice. Note that the cerebellum, which does not express CaMKIIα∷Cre, shows comparable levels of netrin-1 protein in both genotypes. (**c**) Representative traces (left) and group data (right) showing evoked EPSC amplitude in CA1 pyramidal neurons in response to incremental stimulation intensity of Schaffer collaterals from wild-type control littermates (black; Control) and CaMKIIα∷Cre/NTN1^*fl/fl*^ mice (blue; NTN1 cKO). (**d**) Effect of Schaffer collateral high-frequency stimulation (arrow, HFS; 1sX100 Hz) on evoked responses in CA1 pyramidal neurons in acute hippocampal slices from wild-type control littermates (black circles; control, *n* = 9 cells from 6 mice) and CaMKIIα∷Cre/NTN1^*fl/fl*^ mice (blue circles; NTN1 cKO, *n* = 9 cells from 6 mice). (**e**) Representative evoked EPSCs in a CA1 hippocampal neuron in response to Schaffer collateral stimulation before (1 in **d**), and after (2 in **d**) high-frequency stimulation (HFS; 1s of 1×100 Hz) in wild-type control littermates (black) and CaMKIIα∷Cre/NTN1^*fl/fl*^mice (blue). (**f-h**) Group data show AMPAR-mediated current (**f**) (control: 237±30% of baseline, *p*<0.001; NTN1 cKO: 124±9% of baseline, *p*=0.23; 2-way ANOVA: Interaction of genotype X LTP: *F*_1,16_=4.57, *p* = 0.04), NMDAR current (**g**) (control: 106±15% of baseline, NTN1 cKO: 125±24% of baseline; 2-way ANOVA: Main effect of genotype, *F*_1,16_ =0.09, *p* = 0.77; Main effect of simulation, *F*_1,16_=0.02, *p*=0.87; Interaction of genotype X stimulation, *F*_1,16_=1.27, *p*=0.28), and AMPAR-to-NMDAR ratio (**h**) (control: 2.43±0.49 in baseline vs. 4.88±0.48 following HFS, *p*<0.001; NTN1 cKO: 3.16±0.45 in baseline vs. 4.08±0.76 following HFS, *p*=0.09; 2-way ANOVA: Interaction of genotype X stimulation, *F*_1,16_=4.48, *p*=0.049) in wild-type control littermate (*n*=9 cells from 6 mice; black) and CaMKIIα∷Cre/NTN1 −1^*fl/fl*^ mice (*n*=9 cells from 6 mice; blue) before and 20 min after high-frequency stimulation (grey bars).

**Figure 3.**
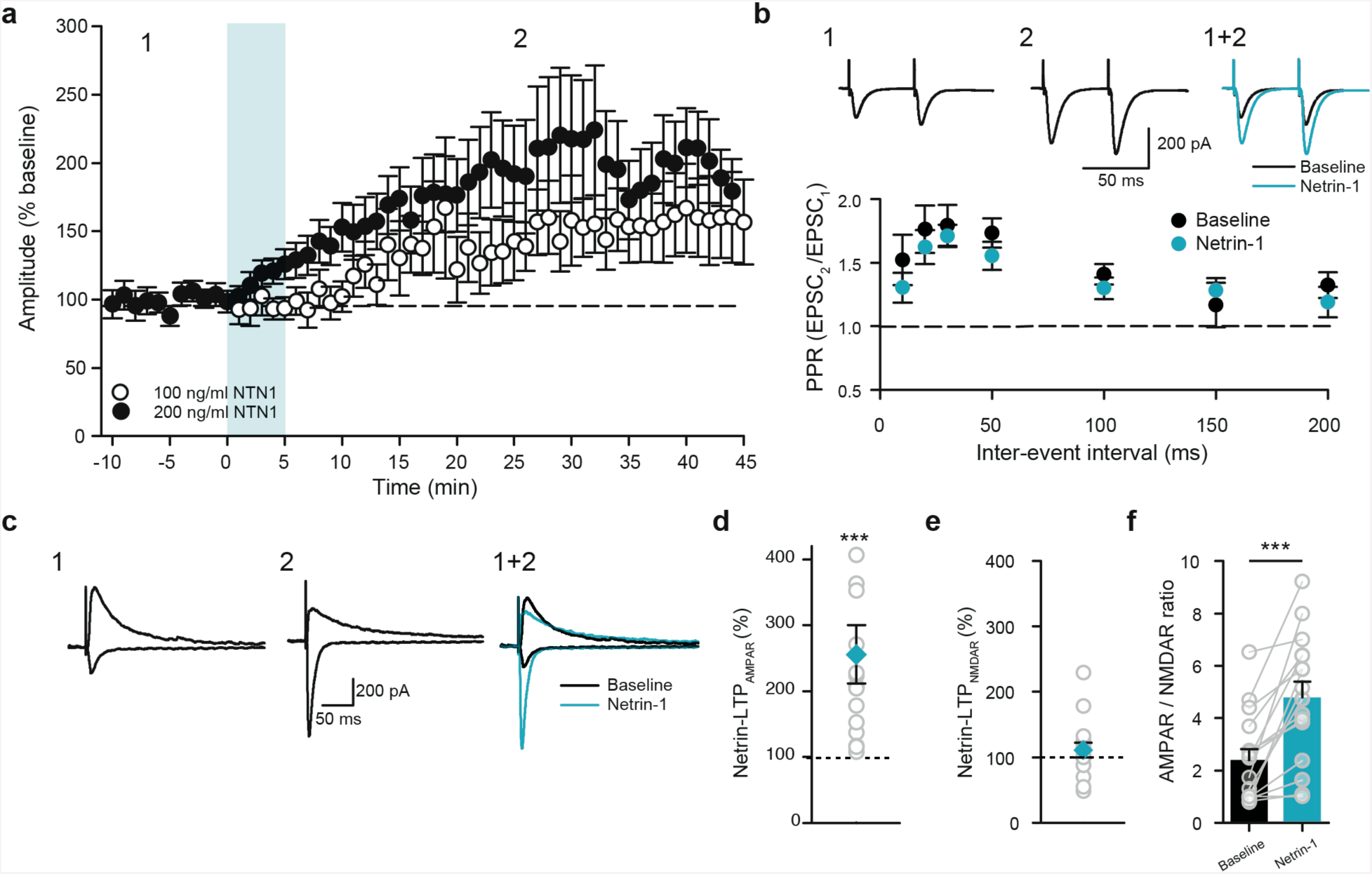
Exogenous netrin-1 potentiates synaptic transmission in adult mouse hippocampus by increasing postsynaptic AMPA receptor-mediated current. (**a**) Effect of 5 min bath application of netrin-1 (blue) on evoked synaptic responses in CA1 pyramidal neurons in response to Schaffer collateral stimulation (100 ng/ml, white circles: *n* = 8 cells from 5 mice; 200 ng/ml, black circles: *n*=16 cells from 10 mice). (**b**) Representative evoked responses (top, 50 ms ISI) in a CA1 pyramidal neuron before (corresponding to 1 in **a**, black trace in overlay) and after (corresponding to 2 in **a**, blue trace in overlay) perfusion with netrin-1 (200 ng/ml). Group data show that paired-pulse ratio is not significantly changed from baseline following 5 min bath application of netrin-1 (200 ng/ml) across a range of ISIs. (Main effect of condition: *F*_1,6_=4.05, *p*=0.09; main effect of ISI: *F*_1,6_=5.22, *p*<0.001; Interaction between condition X ISI: *F*_6,36_=0.35, *p*=0.91; *n*=7 cells from 5 mice; 10 ms, 1.52±0.19 in baseline vs. 1.30±0.12 in netrin-1; 20 ms, 1.76±0.19 in baseline vs. 1.62±0.13 in netrin-1; 30 ms, 1.79±0.16 in baseline vs. 1.71±0.09 in netrin-1; 50 ms, 1.73±0.12 in baseline vs. 1.55±0.11 in netrin-1; 100 ms, 1.41±0.08 in baseline vs. 1.30±0.08 in netrin-1; 150 ms, 1.17±0.18 in baseline vs. 1.28±0.09 in netrin-1; 200 ms, 1.32±0.10 in baseline vs. 1.19±0.12 in netrin-1). (**c**) Representative evoked synaptic responses during baseline (left, 1 in **a**, black trace in overlay) and after netrin-1 (middle, 2 in **a**, blue trace in overlay) at −70 mV (bottom) and +40 mV (top). (**d-e**) Group data show that the AMPAR-mediated current (**d**; 255±44% of baseline, *n*=16, *p*<0.001, Wilcoxon Signed Rank) is significantly increased following bath application of netrin-1 (200 ng/ml), while the NMDAR-mediated current remains unchanged (**e**; 110±11% of baseline, *p* = 0.762). (**f**) Group data show that the AMPAR-to-NMDAR ratio is significantly increased 20 min after bath application of netrin-1 (2.40±0.42 in baseline vs. 4.79±0.61 following netrin-1, *t*_15_=4.99, *p*<0.001, paired-samples *t*-test).

### Netrin-1 is necessary for long-term potentiation at Schaffer collateral synapses

High-frequency stimulation of the Schaffer collaterals induces NMDAR-dependent LTP in the adult hippocampus ^26^. Since NMDAR activation by cLTP stimulation increases NTN1-SEP exocytosis at synaptic sites, we hypothesized that the endogenous release of netrin-1 may contribute to LTP induced by high-frequency stimulation of the Schaffer collateral hippocampal pathway. To test this, we selectively deleted a floxed *ntn1* allele from excitatory neurons by *Cre* expression regulated by the CaMKIIα promoter (T29-CaMKIIα∷Cre/NTN1^*fl/fl*^). *Cre* recombinase is first expressed at ∼2.5 weeks postnatally in these mice, and is detected throughout all subfields of the hippocampus, exclusively in glutamatergic neurons, by one month of age. Critically, expression occurs after developmental axon guidance is complete ^15, 27^. Hippocampal homogenates from 3-6 month old T29-CaMKIIα∷Cre-NTN1^*fl/fl*^ (NTN1 cKO) mice showed reduced levels of netrin-1 protein compared to control littermates (Cre-negative NTN1^*fl/fl*^), consistent with deletion of *ntn1* expression from excitatory forebrain neurons (Figure 2b). Further, immunohistochemical analyses demonstrated that netrin-1 immunoreactivity is substantially reduced in stratum radiatum of NTN1 cKO mice compared to control littermates (Supplemental figure 3).

Previous work has suggested that DCC may function as a dependence receptor, and therefore reducing the level of netrin-1 might increase apoptosis ^28^. However, levels of cleaved caspase-3 were not found to be significantly different between 3-6 month old NTN1 cKO and control littermates (Supplemental figure 3).

To determine if netrin-1 contributes to basal synaptic transmission, we measured evoked excitatory postsynaptic currents (eEPSCs) from CA1 pyramidal neurons following Schaffer collateral stimulation of acute hippocampal slices from 3-6 month old NTN1 cKO and control littermates in the presence of picrotoxin (PTX; 100 µM) to block GABA_A_-mediated synaptic transmission. We detected no significant differences in AMPAR or NMDAR eEPSC amplitudes, paired pulse ratios, rectification, and AMPAR-to-NMDAR ratios indicating that basal evoked synaptic transmission was not affected by deletion of *ntn1* (Supplemental figure 3). To assess whether conditional deletion of netrin-1 impaired the relative strength of individual synapses, we recorded AMPAR-mediated miniature EPSCs (mEPSCs) in CA1 pyramidal neurons from control and NTN1 cKO brain slices in the presence of PTX (100 µM) and tetrodotoxin (TTX; 1 µM). The amplitude of mEPSCs was slightly but significantly reduced in CA1 pyramidal neurons from NTN1 cKO compared to littermate controls, whereas no changes were detected in mEPSC event or instantaneous frequency (Supplemental figure 4). These results provide evidence that conditional deletion of *ntn1* from principal excitatory neurons lead to a modest weakening of synapses onto CA1 pyramidal neurons.

To assess whether netrin-1 may contribute to NMDAR-dependent LTP, we recorded eEP-SCs in CA1 pyramidal neurons from adult NTN1 cKO and control littermates in the presence of the GABA_A_ receptor antagonist, picrotoxin (100 µM). Brief high-frequency stimulation (HFS; 1-s at 100 Hz) resulted in significant short-term facilitation of eEPSCs in CA1 pyramidal neurons from both NTN1 cKO and control littermates. The amplitude of eEPSCs remained significantly potentiated in slices from control littermates after 20-25 min compared to baseline values. However, the amplitude of synaptic potentiation was significantly attenuated in slices from NTN1 cKO mice (Figure 2d-e). Moreover, we observed significant increases in AMPAR-mediated current and AMPAR-to-NMDAR ratio in slices from control littermates following tetanic stimulation, but not in slices from NTN1 cKO mice (Figure 2f-h). These findings demonstrate that deletion of *ntn1* from principal excitatory neurons significantly attenuates NMDAR-dependent LTP, and indicate that netrin-1 plays a central role in synaptic plasticity in the adult hippocampus.

### Netrin-1 potentiates CA3-CA1 Schaffer collateral synapses via an NMDAR-independent postsynaptic mechanism

Genetic studies in *C. elegans* provided initial evidence that netrin directs the local enrichment of synaptic proteins during synaptogenesis ^12, 13, 29^. In mammalian neurons, synaptic proteins, such as PSD95, are rapidly recruited to synaptic sites in response to a local source of netrin-1 ^7^. The impaired LTP detected in T-29-CaMKIIα∷Cre/NTN1^*fl/fl*^ mice suggested that netrin-1 secreted by neurons may be a critical effector of activity-dependent LTP in the adult hippocampus that functions in parallel to or downstream of NMDAR activation. To determine if netrin-1 might be sufficient by itself to elicit changes in synaptic strength, we assessed netrin-1 gain-of-function by perfusing hippocampal slices from adult wild-type C57/B6 mice (2-3 month old) with exogenous recombinant netrin-1. Schaffer collateral-evoked AMPAR-mediated eEPSCs were measured using whole-cell patch clamp recordings from CA1 pyramidal neurons held at −70 mV in the presence of PTX (100 µM). Bath perfusion of netrin-1 (100-200 ng/ml, 5 min) without tetanic stimulation elicited a dose-dependent potentiation of synaptic responses that persisted for at least 40 min (Figure 3a).

To determine if netrin-1 potentiation of eEPSCs was associated with facilitation of presynaptic release, we assessed the paired-pulse ratio across a range of inter-stimulus intervals (10-200 ms ISI) prior to and 20 min after netrin-1 application. We detected no change, consistent with the lack of an effect on presynaptic release properties (Figure 3b). Further, individual paired comparisons failed to detect any change in paired-pulse ratio. These findings suggest that exogenous netrin-1 facilitates AMPAR-mediated synaptic currents in CA1 pyramidal neurons independently of changes in presynaptic glutamate release.

Glutamatergic synapses can be potentiated by the postsynaptic recruitment of AMPARs ^1^. We found that AMPAR-mediated current (measured at −70 mV) was significantly increased following perfusion of netrin-1, whereas NMDAR-mediated current (measured at +40 mV, 50 ms post-tetanus) was unchanged, resulting in an increased AMPAR-to-NMDAR current ratio (Figure 3c-f, Supplemental figure 5). These findings provide evidence that the addition of netrin-1 is sufficient to rapidly and selectively facilitate glutamatergic synaptic transmission in the hippocampus via the postsynaptic insertion of AMPARs into hippocampal synapses.

AMPAR insertion into immature spines can lead to synaptic maturation and contribute to potentiation of synaptic responses in the hippocampus ^30, 31^. To determine if bath application of netrin-1 increases the number of functional synapses by recruiting immature synapses, we recorded AMPAR-mediated mEPSCs in CA1 pyramidal neurons from adult mouse acute hippocampal slices in the presence of PTX (100 µM) and TTX (1 µM) (Figure 4a). Bath application of netrin-1 (200 ng/ml, 5 min) significantly increased mEPSC frequency but not amplitude, consistent with insertion of AMPARs into nascent synapses (Figure 4b-c). To directly assess whether netrin-1 leads to synaptic maturation through the insertion of AMPARs, we recorded evoked AMPAR-mediated currents in response to minimal stimulation. During baseline conditions, low levels of stimulation frequently failed to elicit an evoked inward synaptic current (Figure 4d). Following bath application of netrin-1, the evoked EPSC failure rate was significantly reduced, consistent with an increase in the number of functional glutamatergic synapses recruited (Figure 4e-f). These results provide evidence that netrin-1 recruits AMPARs into immature synapses in adult hippocampal networks.

**Figure 4.**
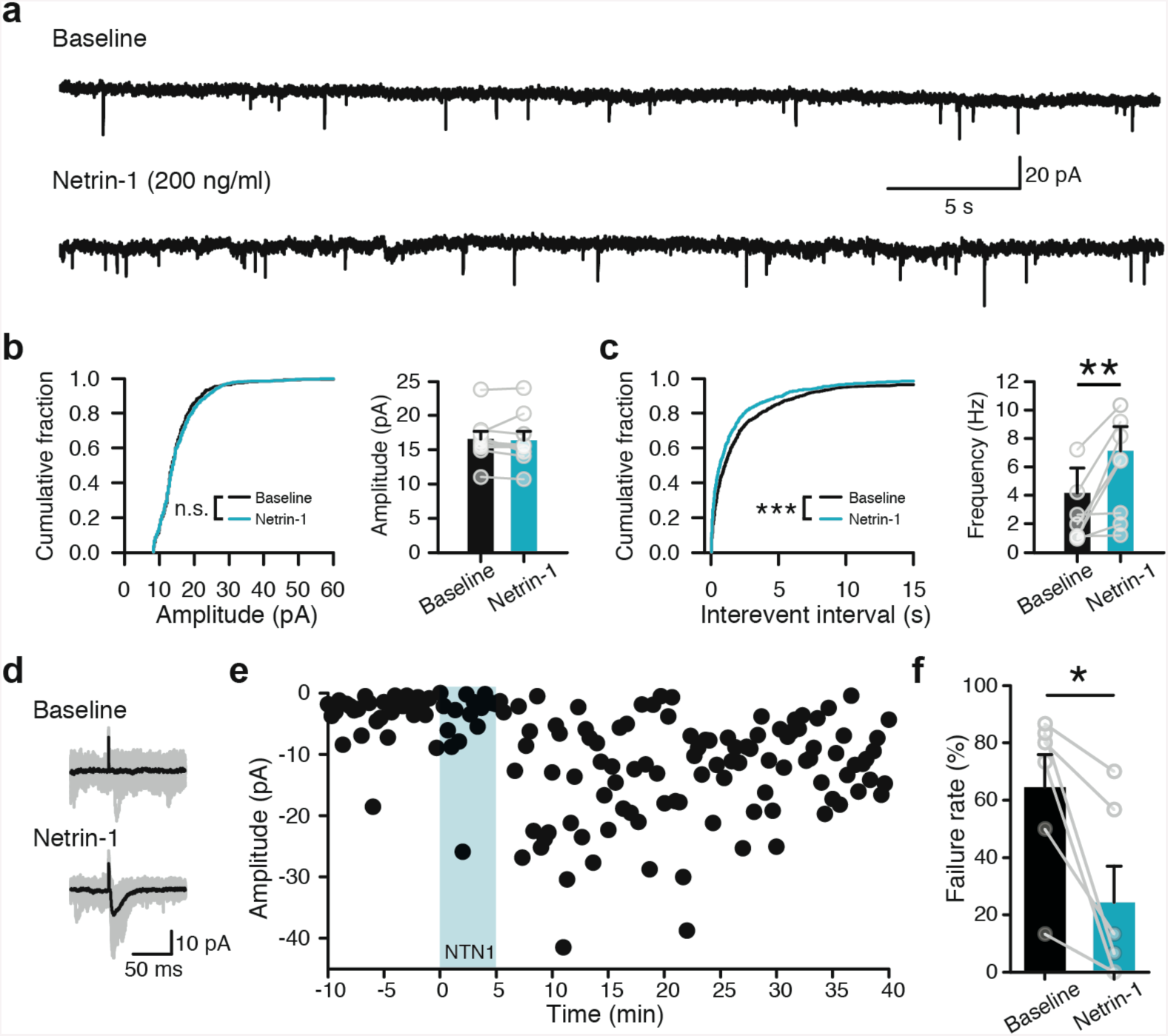
Netrin-1 recruits new synapses in CA1 pyramidal neurons during activity-dependent long-term potentiation. (**a**) Representative voltage-clamp recordings of mEPSCs from CA1 pyramidal neurons held at −70 mV in an acute adult mouse hippocampal slice in the presence of tetrodotoxin (1 µM) and picrotoxin (100 µM) before and after 5 min bath application of netrin-1 (200 ng/ml) (*n*=9 cells from 5 mice). (**b-c**) Cumulative distribution plots (left) and group data (right) of mEPSC amplitude (**b**; 16.5±1.1 pA in baseline vs. 16.4±1.3 pA following netrin-1, *t*_8_=0.35, *p*=0.73, paired samples *t*-test) and frequency (**c**; 4.2±1.8 Hz in baseline vs. 7.1±1.7 Hz following netrin-1, *t*_8_=3.37, *p*<0.01, paired samples *t*-test). Kolmogorov-Smirnov test for cumulative distribution data. (**d-f**) Synaptic responses evoked by minimal stimulation (raw traces in grey, averaged in black, **d**) and individual response amplitudes **(e)** during baseline and 20 min following netrin-1 (blue in **e**). Group data show a significant decrease in failure rate following bath application of netrin-1 (200 ng/ml). (**f**; 64±11% in baseline vs. 24±11% following netrin-1, *t*_5_=3.46, *p*=0.018, paired samples *t*-test; *n*=6 cells from 6 mice).

### Exogenous netrin-1 occludes activity-induced LTP

The rapid and long-lasting potentiation of synaptic responses after bath application of netrin-1, as well as the attenuation of potentiation induced by high-frequency stimulation in hippocampal slices from in T-29-CaMKIIα∷Cre/NTN1^*fl/fl*^mice, suggest that activity-dependent secretion of netrin-1 may contribute to the expression of LTP induced by high-frequency stimulation. To assess whether netrin-1 is linked mechanistically with high-frequency stimulation induced-LTP, we first examined the effect of netrin-1 on previously-potentiated synapses. Following a 5 min baseline period, LTP was induced by brief HFS (1-s at 100 Hz). Synaptic responses were significantly potentiated compared to baseline values and resulted in a significant increase in the AMPAR-to-NMDAR ratio, consistent with insertion of AMPARs at Schaffer collateral synapses in the adult mouse brain following HFS. Bath application of netrin-1 failed to elicit additional enhancement of AMPAR-mediated evoked synaptic responses, nor alter AMPAR-to-NMDAR ratios compared to post-HFS levels (Figure 5a-e). These data indicate that activity-dependent HFS LTP can successfully occlude netrin-1-mediated potentiation.

**Figure 5.**
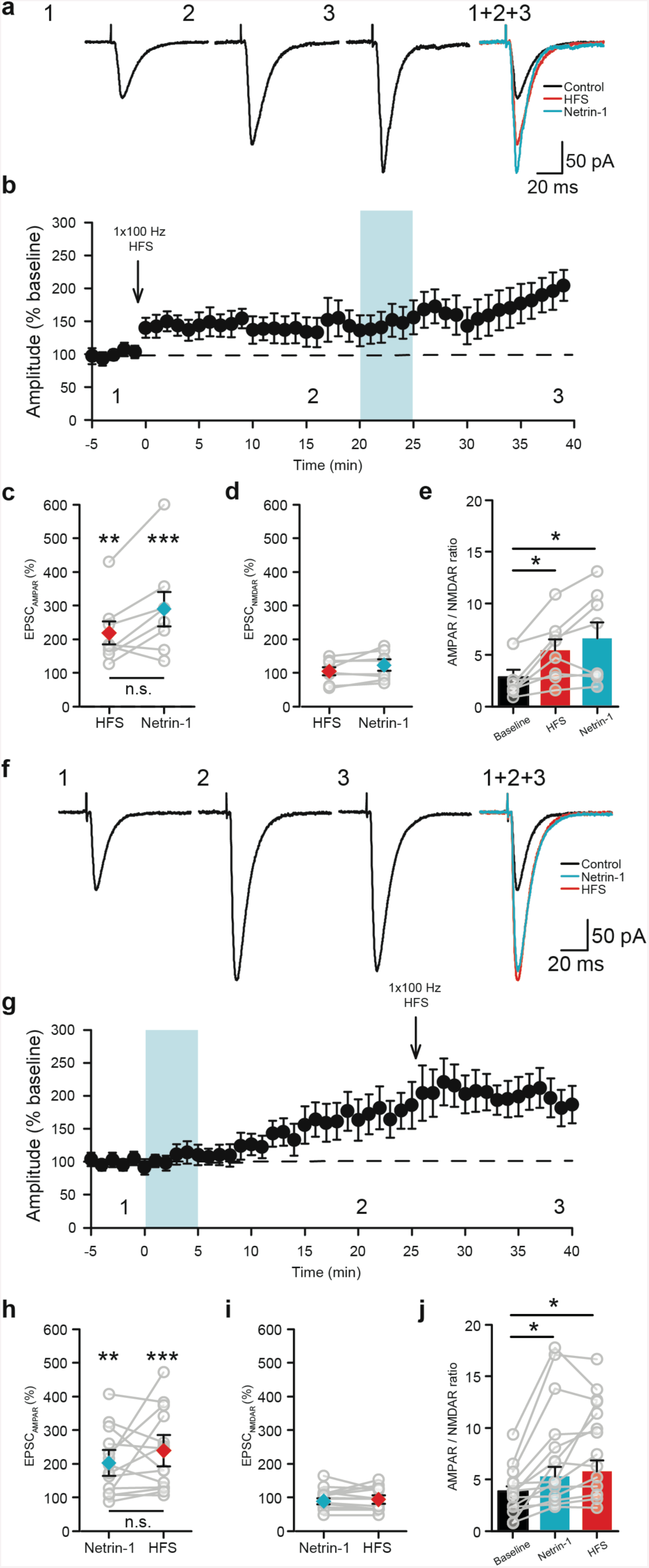
Netrin-1 occludes HFS LTP in the adult hippocampus. (**a**) Representative traces of EP-SCs during baseline (1, black), 20 min following HFS (2, red, 1-s at 100 Hz arrow in **b**), and following 5 min bath application of netrin-1 (200 ng/ml) (3, blue). (**b-e**) Group data show that HFS results in potentiation of AMPAR-mediated synaptic responses (**b**), but does not occlude further potentiation by 5 min netrin-1 application (**c**; 218±34% of baseline following HFS vs. 290±51% following netrin-1, *F*_2,23_=21.84, *p*<0.001, One-way RM-ANOVA; pairwise comparisons, baseline vs. HFS: *p*=0.001, baseline vs. netrin-1: *p*=0.002, HFS vs. netrin-1: *p*=0.106). NMDAR-mediated EPSCs were not significantly changed between all conditions (**d**; 105±12% of baseline following HFS vs. 120±15% following netrin-1, *F*_2,23_=1.05, *p*=0.374, One-way RM-ANOVA), whereas AMPAR-to-NMDAR ratios were significantly elevated following both HFS and netrin-1 (**e**; Baseline: 2.8±0.7, HFS: 5.4±1.0, netrin-1: 6.6±1.5, *F*_2,23_=11.01, *p*=0.006, One-way RM-ANOVA; pair-wise comparisons, baseline vs. HFS: *p*=0.018, baseline vs. netrin-1: *p*=0.029, HFS vs. netrin-1: *p*=0.311). *n*=8 cells from 5 mice. (**f**) Representative traces of evoked synaptic responses during baseline (1, black), 25 min following bath application of netrin-1 (2, blue, 5 min at 200 ng/ml), and following HFS (3, red, 1-s at 100 Hz arrow in **g**). (**g-j**) Group data show that bath application of netrin-1 (blue in **g**) leads to significant potentiation of AMPAR-mediated synaptic responses, but that HFS (1-s at 100 Hz, arrow in **g**) does not further increase evoked amplitude (**h**; 201±26% of baseline following netrin-1 vs. 237±31% of baseline following HFS, *F*_2,13_=11.33, *p*<0.001, One-way RM-ANOVA; pairwise comparisons, baseline vs. netrin-1: *p*=0.001, baseline vs. HFS: *p*<0.001, netrin-1 vs. HFS: *p*=0.92, *n*=8). NMDAR-mediated synaptic responses were not significantly affected by netrin-1 (99±8% of baseline following netrin-1, 98±8% following HFS, *F*_2,13_=1.57, *p*=0.233), whereas AMPAR-to-NMDAR ratio (**i**) was significantly elevated following netrin-1, but was not further increased following HFS (baseline: 3.9±0.5, netrin-1: 7.5±1.4, HFS: 8.5±1.2; *F*_2,13_=17.28, *p*<0.001; baseline vs. netrin-1: *p*=0.004, baseline vs. HFS: *p*<0.001, netrin-1 vs. HFS: *p*=0.39, pairwise comparisons). *n*=14 cells from 9 mice. All pairwise comparisons using Tukey’s multiple comparison test.

We then assessed whether netrin-1-induced potentiation can occlude the expression of LTP induced by high-frequency stimulation. Bath application of exogenous netrin-1 resulted in rapid potentiation of synaptic responses. Twenty min following the application of netrin-1, we attempted to induce LTP using HFS. A single burst of high-frequency stimulation resulted in a brief increase in eEPSC amplitude, consistent with a post-tetanic short-term facilitation of Schaffer collateral inputs. However, by 15 min after high-frequency stimulation we observed no significant potentiation compared to pre-tetanus eEPSC amplitudes (Figure 5f-j). This occlusion suggests that the application of exogenous netrin-1 saturates a mechanism required for high-frequency stimulation-induced LTP. Failure to induce LTP may also have been due to whole-cell dialysis at the induction time point. We therefore applied HFS 30 min after whole-cell break-in and found that it was sufficient to induce LTP (Supplemental figure 6). Together, these findings support the conclusion that high-frequency stimulation promotes netrin-1 secretion, which in turn activates downstream effectors that recruit AMPARs to hippocampal synapses.

### Netrin-1 facilitates synaptic responses via NMDAR-independent increases in intracellular Ca^2+^ levels and activation of CaMKII

NMDAR activation triggers activity-dependent LTP of AMPAR currents ^21^. To assess the role of NMDARs in netrin-1 potentiation of synaptic responses, we bath applied exogenous netrin-1 in the presence of the NMDAR antagonist, APV (50 µM). Netrin-1 resulted in strong potentiation of evoked AMPAR-mediated current (Figure 6a-c), indicating that NMDAR activation is not required for netrin-1 potentiation of synaptic responses. Taken together with the demonstration that NMDAR-dependent cLTP stimulation increases netrin-1 secretion, we conclude that NMDAR activation promotes the release of netrin-1, and that netrin-1 triggers downstream mechanisms that are sufficient to potentiate synaptic responses independent of NMDAR activation.

**Figure 6.**
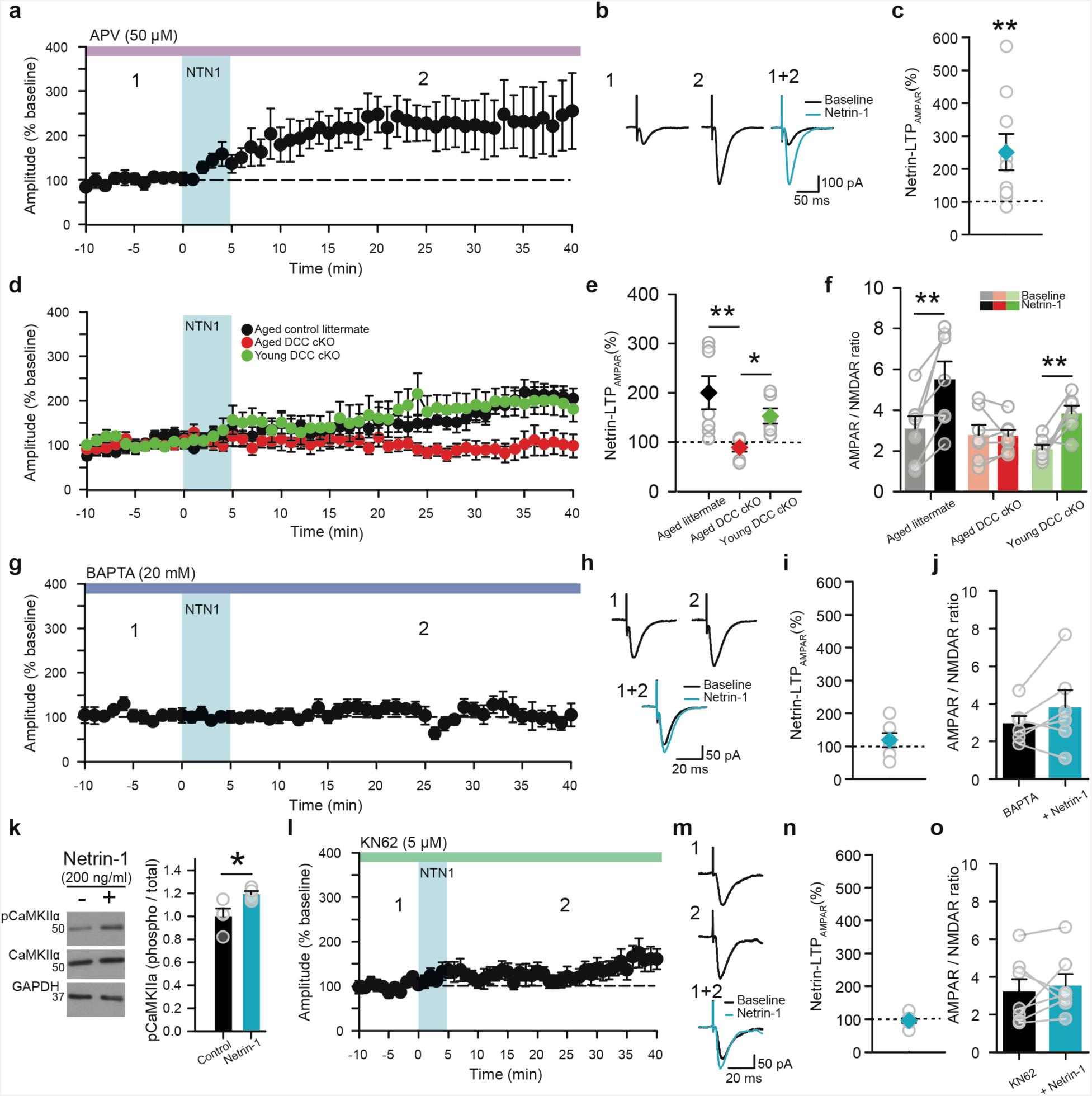
Netrin-1 potentiation of AMPA current requires increased intracellular Ca^2+^ and CaMKII activation independent of NMDAR function. (**a-c**) Evoked synaptic responses in CA1 pyramidal neurons from acute mouse hippocampal slices after bath application of netrin-1 (200 ng/ml, blue) in the presence of APV (**a**; 50 µM, purple; *n*=9 cells from 5 mice). Representative traces (**b**) and group data (**c**) prior to (1 in **a**) and following bath application of netrin-1 (2 in **a**) in the presence of APV (259±53% of baseline; *t*_8_=4.66, *p*=0.001, paired samples *t*-test). (**d-f**) Effect of netrin-1 (**d**; blue bar) on synaptic responses in CA1 pyramidal neurons from acute mouse hip-pocampal slices from aged DCC cKO (>6 months old CaMKIIα∷Cre/DCC^*fl/fl*^, red, *n*=7 cells from 5 mice), young DCC cKO (4-6 weeks old CaMKIIα∷Cre/DCC^*fl/fl*^, green, *n*=5 cells from 3 mice), and control aged littermates mice (>6 months old CaMKIIα∷Cre^*neg*^/DCC^*fl/fl*^, black, *n*=7 cells from 5 mice). (**e**) Group data show that netrin-1 increases amplitude of AMPAR-mediated current in aged littermates and young DCC cKO mice but not in aged DCC cKO (aged control littermate: 200±33% of baseline, *p*=0.002; young DCC cKO: 153±15% of baseline, *p*=0.04; aged DCC cKO: 89±8% of baseline, *p*=0.44; Two-way RM-ANOVA: Interaction between genotype X LTP: *F*_2,17_=5.54, *p* = 0.014) (**f**) AMPAR-to-NMDAR ratio was significantly increased in aged littermates and young DCC cKO following bath application of netrin-1, but not in aged DCC cKO (aged control littermate: 3.0±0.6 in baseline vs. 5.5±0.9 following netrin-1, *p*<0.001; young DCC cKO: 2.1±0.2 in baseline vs. 3.8±0.4 following netrin-1, *p*=0.005; aged DCC cKO: 2.8±0.5 in baseline vs. 2.7±0.3 following netrin-1, p=0.91; Two-way RM-ANOVA Interaction between genotype X LTP: *F*_2,17_=6.22, *p* = 0.009) (**g-i**) Effect of netrin-1 (blue) on average synaptic responses (**g**) and representative evoked EPSCs (**h**) recorded in CA1 pyramidal neurons in acute mouse hippocampal slices with intracellular BAPTA (20 mM, dark blue, *n* = 6 cells from 4 mice). Group data show that netrin-1 did not significantly potentiate AMPAR-mediated synaptic responses (**i**, left; 118±22% of baseline, *t*_5_=0.66, *p*=0.54, paired samples *t*-test) or increase AMPAR-to-NMDAR ratio (**j**; 2.9±0.4 in baseline vs. 3.8±0.9 following netrin-1; *t*_5_=1.47 *p*=0.20, paired samples *t*-test). (**k-o**) Netrin-1 phosphorylates CaMKII to potentiate synaptic responses in the adult hippocampus. (**k**) Representative Western blot (left) and group data (right) from isolated hippocampal slices showing increased levels of phosphorylated CaMKIIα (Thr^286^) following bath application of netrin-1 (200 ng/ml) (control: 1.00±0.06, netrin-1: 1.19±0.02; *t*_6_=2.60, *p*=0.04, independent samples *t*-test, *n*=4 slices per mouse, *N*=4 mice). Effect of netrin-1 (blue bar) on averaged (**l**) and representative (**m**) evoked synaptic responses in CA1 pyramidal neurons in the presence of KN62 (5 µM, *n* =7 cells from 5 mice, green bar). Group data show no significant increase in AMPAR-mediated current (**n**; 95±7% of baseline, *t*_6_=0.73, *p*=0.49, paired samples t-test) or AMPAR-to-NMDAR ratio (**o**, 3.2±0.6 in baseline vs. 3.5±0.6 following netrin-1, *t*_6_=0.56, *p*=0.59, paired samples *t*-test) following netrin-1 (200 ng/ml, 5 min, blue bar).

Netrin-1 binding to DCC mediates axonal chemoattraction during development ^10, 32^. To determine if netrin-1 potentiation requires DCC, we deleted a floxed *dcc* allele from glutamatergic neurons in the adult forebrain by generating T-29-CaMKIIα∷Cre/DCC^*fl/fl*^ mice (DCC cKO) ^15^. Bath perfusion of netrin-1 potentiated eEPSCs in acute slices from control littermates and young 4-6 week old DCC cKO mice that still express high levels of residual DCC, but failed to potentiate eEPSCs in CA1 pyramidal neurons from 6-month-old adult DCC cKO mice (Figure 6d-e). Additionally, while bath application of netrin-1 increased AMPAR-mediated current and AMPAR-to-NMDAR ratios in control littermates and 4-6 week old DCC cKO mice, no increase was detected in aged DCC cKO (Figure 6f). We conclude that netrin-1 potentiation of Schaffer collateral inputs requires DCC.

Both NMDAR and DCC activation result in Ca^2+^ influx and increased intracellular Ca^2+^ concentration, which in turn can promote changes in AMPAR trafficking via Ca^2+^/calmodulin kinase II (CaMKII) activation ^33-35^. To assess the contribution of increased intracellular Ca^2+^ to netrin-1 potentiation, we included the Ca^2+^ chelator 1,2-bis-(2-aminophenoxy)ethane-N,N,N’,N’-tetraacetic acid (BAPTA; 20 mM) in the intracellular recording pipette. BAPTA completely blocked netrin-1 potentiation of AMPAR-mediated synaptic responses and the increase in AMPAR-to-NMDAR ratio (Figure 6g-j), indicating that intracellular Ca^2+^ is critical for netrin-1 potentiation.

CaMKII activation and stargazin phosphorylation are required to trap and accumulate GluA1-containing AMPARs at synapses ^33^. Bath application of netrin-1 resulted in increased CaMKII phosphorylation in hippocampal slice homogenates (Figure 6l), suggesting that netrin-1-mediated activation of CaMKII may potentiate synaptic responses. Consistent with this, bath application of the CaMKII inhibitor KN62 (5 µM) completely blocked netrin-1-induced increases in the amplitude of evoked AMPAR-mediated currents and AMPAR-to-NMDAR ratio in acute hippocampal slices (Figure 6l-o). These results support the conclusion that netrin-1 activates CaMKII, and that CaMKII activation is required for netrin-1 potentiation of evoked responses.

### Netrin-1 potentiation is mediated via postsynaptic accumulation of GluA1-containing receptors

Increases in intracellular Ca^2+^ can trigger AMPAR recruitment to synapses ^34^. AMPARs are composed of homo-or heteromeric configurations of four different subunits (GluA1-4), with Q/R-edited GluA2-containing receptors lacking Ca^2+^ permeability ^36, 37^. Early stages of LTP require interactions mediated by the “TGL” PDZ domain binding motif at the carboxyl terminus of the GluA1 AMPAR subunit ^34^. However, recent studies suggest that multiple forms of AMPARs contribute to this type of synaptic plasticity ^38^. We assessed the subunit composition of AMPARs inserted into the plasma membrane 20 min following application of netrin-1 (200 ng/ml, 5 min) using cell-surface biotinylation of isolated acute adult hippocampal slices. Bath application of netrin-1 significantly increased the amount of cell surface GluA1 compared to control slices from the same animal that were not exposed to netrin-1 (Figure 7a). In contrast, levels of biotinylated GluA2 were not different between conditions (Figure 7b). These findings indicate that netrin-1 selectively increases the amount of GluA1-containing, GluA2-lacking AMPARs at the neuronal surface.

**Figure 7.**
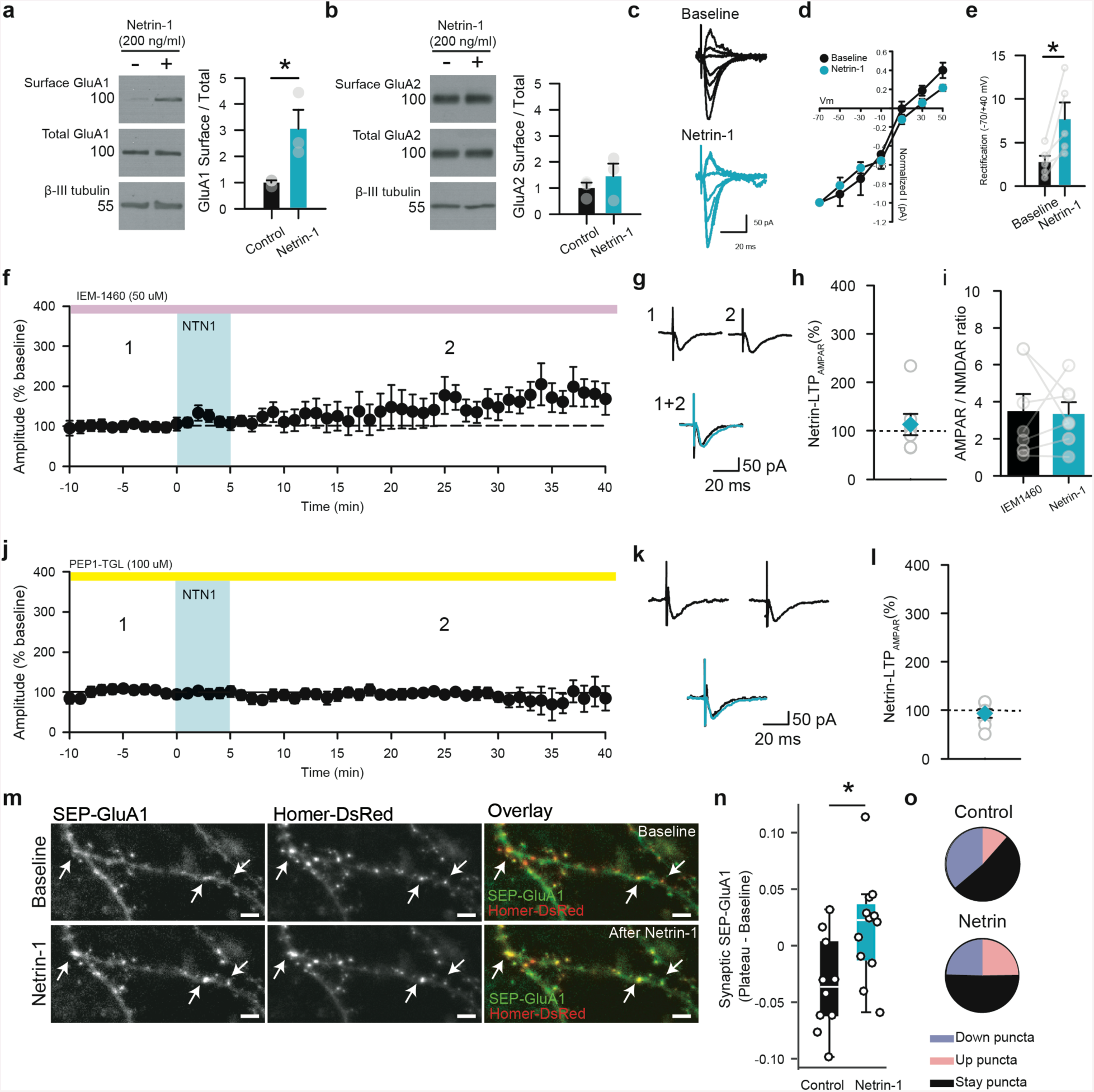
Synaptic recruitment of Ca^2+^-permeable GluA1-containing AMPA receptors by netrin-1. (**a-b**) Western blots (left) and group data (right) of GluA1 **(a**; 306±73% of baseline, *t*_2_=2.81, *p*=0.04, independent samples *t*-test) and GluA2 **(b**; 145±49% of baseline, *t*_2_=0.45, *p*=0.44, independent samples *t*-test) surface membrane distribution in acute hippocampal slices from adult mice with (blue) or without (black) 5 min bath application of netrin-1 (200 ng/ml; *n*=4 slices per mouse, *N* = 3 mice per condition). (**c-e**) Representative traces of synaptic rectification experiments during baseline (black) and following netrin-1 (200 ng/ml; blue). Effect of netrin-1 on I-V curve of AMPAR-mediated synaptic responses (**d**) and rectification index (**e**; 2.7±0.7 in baseline vs. 7.7±1.9 following netrin-1, *t*_4_=3.13, *p*=0.03, paired samples *t*-test). *n* = 5 cells from 3 mice. (**f-i**) Effect of netrin-1 (200 ng/ml; blue) on Schaffer collateral evoked synaptic response in CA1 pyramidal neurons from acute adult hippocampal slices during continuous bath application of the CP-AMPAR antagonist, IEM1460 (**f**; 50 µM, purple bar; *n* = 7 cells from 4 mice). Representative evoked EPSCs (**g**) and group data show AMPAR-mediated current (**h**; 112±22% of baseline, *t*_6_=0.01, *p*=0.99, paired samples *t-*test) and AMPAR-to-NMDAR ratio (**i**; 3.5±0.9 in baseline vs. 3.3±0.9 following netrin-1, *t*_6_=0.16, *p* = 0.87, paired samples *t-*test) in the presence of IEM1460 following bath application of netrin-1. (**j-l**) Inhibition of GluA1 insertion with intracellular Pep1-TGL (100 µM, yellow bar, *n*=8 cells from 4 mice) blocks netrin-1 potentiation of synaptic responses (**j**). Representative evoked EPSCs (**k**) and group data (**l**) show mean AMPAR-mediated current during baseline and following netrin-1 in neurons loaded with Pep1-TGL (**i**; 93±9% of baseline, *t*_7_=0.13, *p*=0.89, paired samples *t*-test). (**m**) Sample images of dendritic segments expressing both SEP-GluA1 (left, green on right) and Homer1C-DsRed (middle, red on right) during baseline (top) and after netrin-1 (200 ng/ml, bottom) in dissociated rat hippocampal neurons. Overlay (right) shows SEP-GluA1 insertion at Homer1C-DsRed sites (white arrows). Scale = 5 µm. (**n**) Average SEP-GluA1 fluorescence variation at Homer1C-DsRed puncta for each neuron (control: ΔF_median_=-0.0364, *n*=10 cells vs. netrin-1: ΔF_median_=0.0226, *n*=12 cells). Wilcoxon test for paired observations, *: *p*<0.05. (**o**) Circular plots show proportion of synaptic states (blue: “down”, black: “stay”, red: “up”; see Methods) after netrin-1 compared to control sister cultures.

GluA2-lacking AMPARs composed of hetero- or homomeric combinations of GluA1, GluA3, and GluA4 subunits are permeable to Ca^2+^, and characterized by pronounced inward rectification due to blockade by endogenous polyamines ^39, 40^. Bath application of netrin-1 in the presence of GABA_A_ and NMDAR antagonists significantly enhanced rectification of AMPAR-mediated currents at depolarized voltages (Figure 7c-e). To determine if netrin-1-induced potentiation is mediated by Ca^2+^-permeable AMPARs (CP-AMPARs), netrin-1 was co-applied with the CP-AMPAR blocker, IEM1460 (50 µM) ^37^. Blockade of CP-AMPARs eliminated the netrin-1 induced potentiation of synaptic responses, and blocked the increases in AMPAR-mediated current and AMPAR-to-NMDAR ratio (Figure 7f-i). These findings are consistent with GluA2-lacking CP-AMPARs underlying the netrin-1-induced synaptic potentiation, likely engaging the incorporation of hetero- or homotetramers of GluA1. To test this, we then blocked the incorporation of GluA1-containing AMPARs by including the competitive peptide Pep1-TGL (100 µM) in the intracellular solution ^34, 41^. Pep1-TGL, which prevents SAP97 binding to the “TGL” PDZ binding motif at the GluA1 C-terminus, completely blocked netrin-1 potentiation of AMPAR-mediated current (Figure 7j-l). To determine if netrin-1 promotes GluA1 synaptic localization, we co-expressed super ecliptic pHluorin-tagged GluA1 (SEP-GluA1) and the post-synaptic marker Homer1c-DsRed in cultured rat hippocampal neurons (14 DIV). Bath application of netrin-1 (200 ng/ml) triggered a rapid increase in SEP-GluA1 fluorescence at a subset of synapses compared to cultures not treated with netrin-1 (Figure 7m-n, Supplemental movie 2-3). Moreover, bath application of netrin-1 resulted in an increase in the number of SEP-GluA1 positive synapses (“up” puncta), while decreasing the number of SEP-GluA1 negative synapses (“down” puncta) compared to control cultures without netrin-1 application (Figure 7o). Together, these findings indicate that netrin-1 promotes the insertion of GluA1-containing AMPARs at a subset of synaptic sites in hippocampal pyramidal neurons.

## Discussion

Netrin-1 is essential for normal embryonic neural development, directing cell migration, axon guidance, and synaptogenesis. Neuronal expression and enrichment at synapses in the adult CNS suggested that netrin-1 might also influence the plasticity of mature neural circuits. Here, we demonstrate that netrin-1 expressed by neurons plays a critical role in the modification of glutamatergic transmission at Schaffer collateral synapses between CA3 and CA1 pyramidal neurons in the adult mammalian hippocampus. Our findings reveal that netrin-1 is secreted by neurons in response to depolarization and NMDAR activation. We demonstrate that neuronal expression of netrin-1 in principal forebrain neurons is critical for activity-dependent LTP, that brief application of exogenous netrin-1 triggers long-lasting synaptic potentiation, and that netrin-1-induced synaptic potentiation occludes LTP induced by high-frequency stimulation. Further, we show that netrin-1-induced potentiation enhances glutamatergic synaptic transmission through DCC-mediated recruitment of GluA1-containing AMPARs to hippocampal synapses in a CaMKII-dependent manner. These findings reveal an essential role for neuronal secretion of netrin-1 in glutamatergic synaptic plasticity in the adult hippocampus that is critical for classic activity-dependent LTP.

### Netrin-1 contributes to activity-dependent LTP

Long-lasting changes in synaptic strength in the adult hippocampus depend on activation of postsynaptic NMDARs ^42^. Here, we report that netrin-1 is enriched in dendrites (Figure 2) and that NMDAR activation can result in netrin-1 exocytosis (Figure 1), suggesting a role in activity-dependent synaptic plasticity in the hippocampus. Consistent with this, conditional loss of netrin-1 from excitatory hippocampal neurons results in attenuated expression of LTP with no change in short-term plasticity following HFS (Figure 2). Short-term facilitation of synaptic responses following HFS is mediated through changes in presynaptic Ca^2+^ that enhance transmitter exocytosis ^43^. In contrast, changes in the distribution of post-synaptic receptors can be detected within 5-10 min after HFS, consistent with the time course observed following exogenous application of netrin-1. Moreover, our findings indicate that following LTP-inducing stimulation, the NMDAR-mediated Ca^2+^ influx mobilizes the secretion of netrin-1 from dendrites at or near synaptic sites (Figure 1). We propose that netrin-1 then activates postsynaptic signaling mechanisms such as CaMKII, independent of NMDAR activation, that locally recruit GluA1-containing CP-AMPARs to excitatory synapses, resulting in postsynaptic potentiation of synaptic transmission.

### Activity-dependent secretion of netrin-1 enhances synaptic transmission through maturation of immature synapses

Intracellular vesicles isolated from adult rat brain synaptosomes contain readily detectable netrin-1 and DCC. Patches of DCC immunoreactivity decorate dendritic spines, and DCC is associated with a detergent-resistant component of postsynaptic density ^15^. Postsynaptic expression of LTP requires soluble N-ethyl-maleimide-sensitive factor-attachment proteins (SNAPs), which promote vesicle exocytosis in a t-SNARE-dependent manner ^44, 45^. DCC interacts with the t-SNARE synatxin-1 and v-SNARE TI-VAMP. However, it remains to be demonstrated that netrin-1 activation of DCC in dendritic spines is required for SNARE function during activity-dependent AMPAR recruitment.

Netrin-1 is secreted from Homer1C-positive sites along dendrites in response to plasticity-inducing stimulation (Figure 1). We also observed exocytotic events at sites lacking Homer1C-DsRed. Homer1C is an adaptor for PSD enriched proteins, where it regulates glutamatergic synaptic transmission through interactions with Group I metabotropic glutamate receptors (mGluR1/5) ^46, 47^. Recent evidence indicates that ∼20% of excitatory synapses in the hippocampus may lack the PSD proteins necessary for synaptic stabilization ^48^, and that these non-PSD-containing synapses exhibit relatively low AMPAR to NMDAR ratios ^49, 50^. Bath application of netrin-1 potently increases the AMPAR-to-NMDAR ratio (Figure 3), indicating that netrin-1 secretion promotes insertion of GluA1-containing AMPARs, recruiting these receptors to immature synapses, possibly including those that initially lack PSD proteins. This conclusion is further supported by the demonstration that challenging cultured rat pyramidal neurons with an immobilized netrin-1 coated micro-bead is sufficient to rapidly recruit PSD-95 ^7^, a critical post-synaptic adaptor that drives glutamate receptor clustering and promotes synaptic maturation ^51^.

Previous work suggests that HFS LTP selectively recruits GluA1-containing CP-AMPARs and increases AMPAR single channel conductance ^52, 53^. Consistent with increases in immature synapses in the absence of netrin-1, conditional deletion resulted in significantly decreased amplitude of mEPSCs compared to control littermates, suggesting that lack of netrin-1 results in reduced AMPAR density at synapses (Supplemental figure 4). In contrast, following bath application of exogenous netrin-1 to acute hippocampal slices from wild-type mice, we observed ∼2 fold increase in mEPSC frequency with no appreciable change in mEPSC amplitude. These findings suggest that netrin-1 may recruit of CP-AMPARs to immature synapses to serve as “placeholders” for subsequent synaptic stabilization. Indeed, previous work has suggested that LTP can lead to increases in mEPSC frequency ^54, 55^ by increasing the density of mature synapses ^56, 57^. Therefore, netrin-1 may facilitate the maturation of immature synapses by promoting the incorporation of transient GluA1-containing AMPARs, which may be replaced with GluA2-containing AMPARs by synaptic local protein synthesis ^14, 58, 59^.

AMPAR trafficking is a key process for LTP expression in the hippocampus, and is mediated in part through NMDAR-mediated increases in intracellular Ca^2+^ and CaMKII activation ^60,61^. Interestingly, LTP results in enlargement of thin-type dendritic spines, which are largely devoid of AMPARs under control conditions ^62, 63^. Increases in thin-type spine volume is dependent, in part, on activation of CaMKII and RhoGTPases including RhoA and Cdc42 ^64, 65^. Netrin-1 binding to DCC, which is critical for netrin-1-mediated potentiation of AMPAR currents (Figure 6), directs cytoskeletal reorganization in developing neuronal growth cones by regulating Cdc42, Rac1 and RhoA ^66-68^. However, it remains to be determined how netrin-1 may contribute to AMPAR trafficking and structural modification of dendritic spines associated with LTP in the adult hippocampus.

We have demonstrated that netrin-1 is enriched in the stratum radiatum of adult mice and is detected intracellularly within dendritic shafts and spines of CA1 pyramidal neurons (Figure 2). Our findings suggest that NMDAR-dependent exocytosis of netrin-1 during activity-dependent LTP promotes recruitment of Ca^2+^-permeable GluA1-containing AMPARs at immature or nascent synapses. Although certain forms of activity-dependent plasticity do not require GluA1 receptor insertion ^38^, trafficking of GluA1-containing AMPARs is dependent on increases in intracellular Ca^2+^ and is critical for LTP in the adult hippocampus ^52, 69, 70^. Application of netrin-1 is sufficient to trigger a local increase in the concentration of intracellular Ca^2+^ in developing neurons ^35, 71-73^. We propose that NMDAR-mediated synaptic Ca^2+^ transients in dendritic spines facilitate netrin-1 release, which in turn locally increases intracellular Ca^2+^ from internal stores to activate CaMKII, and promote the focal insertion and synaptic trapping of GluA1-containing AMPARs. Together, our findings indicate that NMDAR-mediated increases in intracellular Ca^2+^ promote the activity-dependent secretion of netrin-1, which functions as a critical effector of long-term synaptic plasticity.

## Acknowledgments

The authors thank Wayne Sossin, and the members of the Kennedy, Ruthazer, Séguéla, and De Koninck labs for comments on drafts of the manuscript. We also thank Francine Nault, Charleen Salesse, Nathalie Marcal, Yi Jiang, and Hanan Elimam for technical assistance. S.D.G. was supported by postdoctoral fellowships from Fonds de la Recherche Québec – Santé (FRQS) and Canadian Institute for Health Research (CIHR). I.V.B. was supported by a graduate scholarship from Natural Sciences and Engineering Research Council (NSERC) of Canada. J.G. was supported by a postdoctoral fellowship from FRQS. The project was supported by grants from NSERC (P.D.K., P.S., E.S.R.), CIHR (R.A.M., P.D.K., P.S., E.S.R., T.E.K.) and the Alzheimer Society of Canada (T.E.K.). E.S.R. holds a FRQS Research Chair.

## Author contributions

S.D.G., S.L., I.V.B, P.D.K., E.S.R., and T.E.K. conceived the study and designed the experiments. S.D.G. and J. G. performed and analyzed electrophysiological data. S. D. G. and I.V.B. performed and collected molecular biological data. S.L. performed live imaging experiments. S.D.G. and S.A. performed and collected immunohistological data. D.H., S.N.H., R.A.M., and P.S. provided reagents and resources. S.D.G, S.L., P.D.K, E.S.R., and T.E.K. wrote the paper, and all authors read and approved the final manuscript. P.W., P.S., P.D.K., E.S.R., and T.E.K supervised the study.

## Methods

### Animals

The study was conducted in accordance with the guidelines of the Canadian Council for Animal Care and approved by the McGill University and Université Laval Animal Care Committees. All animals were housed in group housing, and provided *ad libitum* access to food and water. For experiments using wild-type animals, C57/B6 mice (8-12 weeks) were used (Charles River, St Constant, Canada). Thy1-GFP (line 15) mice were used in immunohistochemical experiments, and express low levels of membrane-targeted EGFP (mGFP)-positive cells within the CA1 region of the hippocampus under the control of Thy1.2 promoter ^74^. Floxed alleles of *ntn1* (NTN1^*fl/fl*^) and *dcc* (DCC^*fl/fl*^) were generated in mice and maintained on a C57BL/6 genetic background as described ^15, 25^. T29-1 CaMKIIα∷Cre mice (JAX 005359) were obtained from The Jackson Laboratory (Bar Harbor, ME, USA) and maintained on a C57BL/6 background. Both male and female T-29-CaMKIIα∷Cre/netrin-1^*fl/fl*^ (T-29-CaMKIIα∷Cre/NTN1^*fl/fl*^ > 3 months old) and T-29-CaMKIIα∷Cre/DCC^*fl/fl*^ (T-29-CaMKIIα∷Cre/DCC^*fl/fl*^ > 6 months old) mice were used for molecular biological and electrophysiology experiments. We observed no statistically significant differences between sexes, and therefore all data were pooled for analysis. Control experiments were performed using littermates that were negative for *Cre* and homozygous for floxed alleles of *ntn1* and *dcc*.

### Hippocampal neuronal culture

For cell biological assays, hippocampi from E18 or P1 Sprague Dawley rats were isolated, dissociated, plated and cultured as described ^7, 15, 75^.

### Immunohistochemistry

Immunohistochemistry was performed on free-floating sections derived from Thy1-GFP (line 15). Briefly, animals were deeply anaesthetized by intraperitoneal injection of a mixture of 2,2,2 – tribromoethyl alcohol and tert-amyl alcohol diluted at 2.5% in PBS, and transcardially perfused with cold PBS (pH: 7.4) followed by 4% paraformaldehyde (PFA). Brains were removed and post-fixed for 24 h in 4% PFA, followed by 24 h cryoprotection in 30% sucrose in PBS. Thirty µm sections were cut using a cryostat (Leica CM1850), and stored in PBS at 4°C.

Antigen retrieval was performed by boiling sections in 0.1M citrate buffer for 10 min, and allowing to cool to RT. Sections were then washed three times in PBS-T (0.3% Triton X-100), and blocked for 1.5 h in PBS-T containing 3% BSA. Brain sections were incubated sequentially overnight with rabbit monoclonal anti-netrin-1 (1:500, Abcam, RRID:AB_11131145) and chicken polyclonal anti-GFP (1:1000, Abcam, RRID:AB_300798), washed 3 ×; 10 min in blocking solution, and coupled to corresponding secondary antibodies (donkey anti-rabbit Alexa 555, 1:1000, RRID:AB_2536182; and goat anti-chicken Alexa 488, 1:1000, RRID:AB_2636803) for 1h at RT. Sections were then successively washed twice for 10 min with PBS-T and PBS, mounted on gelatin-coated glass microscope slides, and coverslipped using DAPI-Fluromount-G (Southern Biotech).

Embryonic hippocampal neurons infected with AAV8-hSyn-HA-hM3D(Gq)-IRES-mCitrine were fixed with 4% PFA, washed 3 ×; 10 min with 0.1% PBS-T, blocked in 0.1% PBS-T containing 3% BSA for 1h, incubated overnight in rabbit anti-HA (1:1000, Abcam, RRID:AB_307019), washed 3 ×; 10 min with 3% BSA in 0.1% PBS-T, and coupled with corresponding secondary anti-rabbit Alexa 488 (1:1000, ThermoFisher Scientific, RRID: AB_143165) and Hoescht stain (1:1000, ThermoFisher Scientific, RRID:AB_2307445). Coverslips were washed 2 ×; 5 min with PBS-T followed by 2 ×; 5 min in PBS and mounted on glass microscope slides.

Immunofluorescence images were obtained using either a Zeiss Axiovert (S100TV) with epifluorescence illumination at 40x (0.75 N.A.), or a Zeiss LSM 880 laser-scanning confocal microscope equipped with an Airyscan detection unit using a high NA oil immersion alpha Plan-Apochromat 100X/1.46 Oil DIC M27 (Zeiss) with a zoom adjustment to 1.8. All imaging was performed using Immersol 518 F immersion media (ne = 1.518 (23 °C); Carl Zeiss). Detector gain set to 700 and pixel dwell times were adjusted for each dataset keeping them at their lowest values in order to avoid saturation and bleaching effect. The pinhole was adjusted to have an AU of ∼1.25 considering the axial position in tissue. The different fluorophores were acquired sequentially with appropriate filter conditions. The information is listed in the following as “fluorophore – excitation wavelength – dichroic mirror – detection filter”. Alexa Fluor 488 – 488 nm (Argon) – MBS488 – BP 495-550 + LP 570; CF 568 – 561 nm (DPSS 561-10) - MBS 458/561 – BP 495-550 + LP 570; Alexa Fluor 647 – 633 nm (HeNe633) – MBS 488/561/633 – BP 570-620 + LP 645. For processing Airyscan images, Zen Black 2.3 software was used to enhance the spatial resolution 1.7-fold of the diffraction limit by processing each of the 32 Airy detector channels separately with filtering, deconvolution and pixel reassignment. The visualization and rendering was processed with Imaris 8.1.2 (Bitplane AG, Zurich CH).

For images in Supplemental figure 2c-e, cultured hippocampal neurons were fixed for 10 min in 4% PFA solution (0.1 M phosphate buffer, 4% sucrose, and 2 mM EGTA, pH 7.4). Neurons were then washed once with PBS and twice with 0.1 M PBS/glycine and were permeabilized for 30 min in blocking solution (PBS, 2% normal goat serum, and 0.1% Triton X-100). Primary antibody against netrin-1 (1:500, Abcam, RRID:AB_11131145) was diluted in blocking solution and incubated for 2 h at room temperature. After washes in PBS, secondary antibodies (goat anti-rabbit STAR488, Abberior, 1:1000) was diluted in blocking solution and incubated for 45 min at room temperature. Coverslips were mounted in Prolong Gold Antifade mounting media (Thermofisher). Images were acquired on a confocal system (LSM700; Carl Zeiss) using a 63x 1.4 NA oil immersion objective.

### Protein media concentration

For protein media concentration experiments, rat hippocampal neurons from E18 Sprague Dawley rats were plated at 2.5 ×; 10^6^ cells and used at 14 DIV. For hM3D/CNO experiments, cells were infected at 6 DIV with pAAV8-hSyn-HA-hM3D(Gq)-IRES-mCitrine (MOI: 1:10000; Neurophotonics Molecular Tools Platform, Université Laval, Québec, QC, Canada). At 14 DIV, cells were washed 3x with warm (37°C) control ACSF containing (in mM): 135 NaCl, 3.5 KCl, 1.2 MgCl_2_, 2 CaCl_2_, 10 HEPES, and 20 D-Glucose (pH: 7.4, 300 mOsm). Cells were then incubated at 37°C in ACSF with or without clozapine-N-oxide (CNO, 10 µM) for 1 h. For KCl experiments (Supplemental figure 1), cells were washed 3x with serum-free warmed NeuroBasal media (NBM, ThermoFisher Scientific). Cells were then incubated in NBM supplemented with high KCl (20 mM) for 20 min at 37°C. Protease inhibitors (in 1 mg/ml aprotonin, 1 mg/ml leupeptin, 100 mM PMSF, and 0.5 M EDTA) were then added to the media, and cells and media were rapidly cooled to 4° C. Equal volumes of supernatant were collected into a 4 ml Amicon Ultra Centrifugal tube with Ultracel 10-membrane (EMD Millipore), and goat anti-chicken Alexa 633-conjugated secondary antibody (40 ng per tube) was added to ensure equal concentration. Control and experimental media tubes were then centrifuged at 5,000 x g for 60 min. The resultant supernatant was concentrated at 200x, lysed in Laemmli sample buffer, and boiled for 5 min. Cells were harvested and processed as described above.

### Cell surface biotinylation and phosphorylation assays in acute brain slice

Cell surface biotinylation and phosphorylation assays were performed as described ^76, 77^. For surface biotinylation in brain slices, acute hippocampal slices from 2-3 month old C57/B6 mice were made using the same protocol as for electrophysiological experiments. Slices were then transferred to a recording chamber and continuously perfused with control ACSF. Exogenous netrin-1 (200 ng/ml) or vehicle was bath applied for 5 min, followed by a 20 min wash in ACSF (ACSF). Individual hippocampi were isolated by microdissection in ice-cold ACSF, and washed 3x in cold ACSF. For biotinylation assays, 3-4 hippocampi from the same animal and condition were transferred to chilled ACSF containing Ez-link Sulfo-NHS-LC-biotin (1 mg/ml; ThermoFisher Scientific) for 45 min. The biotin reaction was then quenched by washing the slices twice for 25 min at 4° C in 10 mM glycine in ACSF, followed by 3x in cold ACSF (5 min). Slices were harvested in RIPA buffer (containing 150 mM NaCl, 20 mM Tris, pH 8.0, 1%, NP-40 (USB Corp., Cleveland, OH, USA), 0.5% sodium deoxycholate, 0.1% SDS, 1 mM EDTA) supplemented with protease inhibitors (1mg/ml aprotonin, 1mg/ml leupeptin, 100 mM PMSF, and 0.5M EDTA). Biotinylated proteins were precipitated using streptavidin-agarose beads (ThermoFisher Scientific) at 4°C for 2h, centrifuged at 13,800 x g for 15 min, and washed twice with cold PBS. Proteins were then eluted from beads using Laemmli sample buffer, and boiled for 5 min.

For phosphorylation assays, hippocampi were dissected in ACSF and harvested in RIPA buffer supplemented with protease and phosphatase inhibitors (1mM Na_3_V0_4_ and 1mM NaF), homogenized, and centrifuged for 10 min at 13,800 x g. Supernatant was then collected, lysed in Laemmli sample buffer, and boiled for 5 min.

### Western blot analysis

Netrin-1 expression was assessed from hippocampal and cerebellar homogenates derived from 2-3 month old C57/B6 or 3 month old T-29-CaMKIIα∷Cre/NTN1^*fl/fl*^ mice. Briefly, mice were deeply anaesthetized, and the brain was quickly extracted in ice-cold Ringer solution. Cerebellum, cortex, and hippocampi were dissected and isolated in cold DMEM (Life Technologies) supplemented with protease and phosphatase inhibitors, and homogenized in cold RIPA buffer containing (10 mM phosphate buffer [pH 7.2], 150 mM NaCl, 1% NP-40, 0.5% sodium deoxycholate, and 0.1% SDS) with protease and phosphatase inhibitors (aprotinin 2 mg/ml, leupeptin 5 mg/ml, EDTA 2 mM, sodium orthovanadate 1 mM, sodium fluoride 1 mM, and PMSF 1 mM).

Extracts from protein media concentration supernatant, cell lysates, whole hippocampal homogenates, or hippocampal slice homogenates were assessed using Bradford protein assay (Pierce BCA kit, ThermoFisher Scientific). Equal protein levels were loaded and subjected to 10% SDS-page gel electrophoresis and transferred to nitrocellulose membranes, as described ^15^. Western blots were analyzed using mouse beta-III tubulin (1:1000; Sigma Aldrich, RRID:AB_477590), rabbit anti-active caspase 3 (1:1000; Abcam, RRID:AB_302962), rabbit anti-CaMKII (1:1000; ThermoFisher, RRID:AB_2533032), mouse anti-DCC (1:1000; BD Pharmingen, RRID:AB_395314), chicken anti-GAPDH (1:5,000; Millipore, RRID:AB_11211911), rabbit anti-GluA1 (1:10,000; Abcam, RRID:AB_10860361), rabbit anti-GluA2 (1:10,000; Abcam, RRID:AB_2620181), rabbit anti-netrin-1 (1:10,000; Abcam, RRID:AB_11131145), rabbit anti-phospho-Thr^286^(1:1000; PhosphoSolutions, RRID: AB_2492051). Blots were developed using Immobilon Western Chemiluminescent HRP Substrate (Millipore). Densitometry and quanti?cation of relative protein levels were performed on scanned images of immunoblots using Fiji image software ^78^.

### Optical imaging of NTN1-SEP

NTN1-SEP was generated by subcloning the SEP coding sequence (GenBank: AY533296.1) ^79^ lacking the initiating methionine in frame at the C terminus of the sequence encoding netrin-1 ^9^ in plasmid eGFP-N3 (Clontech). For optical imaging of NTN1-SEP, rat hippocampal neurons were plated at a density of 75 cells / mm^2^on glass coverslips coated with poly-D-lysine and maintained in Neurobasal/B27 (Thermo Fisher Scientific). Neurons were transfected with plasmids encoding NTN1-SEP and Homer-DsRed using Lipofectamine 2000 (Thermo Fisher Scientific) at 12 DIV for NTN1-SEP imaging, as described ^80^. Neurons (13-14 DIV) were imaged at 32-35 °C in an open perfusion chamber (0.2-0.5 ml/min) (Warner Instruments) mounted on an Olympus IX-71 inverted microscope equipped with a 100X objective (N. A. = 1.49) with Toptica Chrome MLE laser sources, and a backlit thinned CCD (Princeton ProEm 512B-FT). Signals for NTN1-SEP and Homer1C-DsRed were discriminated using the following laser lines and Semrock filters: SEP, excitation 488nm, emission Brightline 520/35; Homer1C-DsRed, excitation at 560 nm, emission with BrightLine 617/73.

To image netrin-1 exocytosis, a segment of dendrite expressing NTN1-SEP and Homer1C-DsRed was selected, and subsequently photobleached to 30-40% of the initial fluorescence by exposing the dendrite to approximately 40s of high power laser (488nm) light. Movies of 1000 images, at 100 ms per image, were then acquired at lower laser power, with neurons in baseline extracellular solution (in mM: 102 NaCl, 5 KCl, 10 Na HEPES, 1 MgCl_2,_ 1.2 CaCl_2_, 10 Glucose, pH 7.3-7.4, 225-230 mOsm) followed by cLTP solution (same as baseline, except lacking Mg^2+^, and with 200 µM glycine and 30 µM bicuculline).

Image analysis was performed using a custom-made Matlab (Mathworks Inc., Natick, USA) script allowing semi-automatic detection of exocytotic events. Based on any single exocytotic event occurring faster than the image integration time, we computed maps of the events by subtracting each image from the previous frame. Each map was then segmented to identify regions exhibiting a fluorescence increase, which were extracted to calculate the ΔF/F on 500 ms preceding the event and 1500 ms following. We then computed the area under the ΔF/F curve, using a detection threshold of 2.

### Brain slice *in vitro* electrophysiology

Acute horizontal brain slices containing the hippocampus were obtained from adult C57/B6 mice (8 to 12 weeks), and CaMKII∷Cre/NTN1^*fl/fl*^ (3-6 months old) and age-matched control littermates. Mice were deeply anaesthetized by intraperitoneal injection of a mixture of 2,2,2 – tribromoethyl alcohol and tert-amyl alcohol diluted at 2.5% in PBS, and transcardially perfused with ice-cold choline chloride-based solution containing (in mM): 110 choline-Cl, 1.25 NaH_2_PO_4_, 25 NaHCO_3_, 7 MgCl_2_, 0.5 CaCl_2_, 2.5 KCl, 7 glucose, 3 pyruvic acid, and 1.3 ascorbic acid, bubbled with carbogen (O_2_ 95%, CO_2_ 5%). The brain was rapidly removed, thick horizontal brain slices (300 µm) containing the hippocampus were cut using a vibrating microtome (VT1000s, Leica), and allowed to recover for 1h in artificial cerebrospinal fluid (ACSF) containing, in mM: 124 NaCl, 5 KCl, 1.25 NaH2PO4, 2 MgSO4, 26 NaHCO3, 2 CaCl, and 10 Glucose saturated with 95% O2 and 5% CO2 (pH ∼7.3, 300 mOsm) at room temperature (22°-24° C).

Individual brain slices were placed in a custom-built recording chamber, and continuously perfused with warmed (30 ± 2° C) ACSF (TC324B, Warner Instruments). Voltage-recordings of CA1 hippocampal pyramidal neurons were performed on an upright microscope (Nikon Eclipse or Scientifica SliceScope 2000) equipped with a micromanipulator (Sutter MP-225 or Scientifica Patchstar), a 40x or 60x water immersion objective (0.8 or 1.0 N.A., respectively), differential interference contrast optics, and coupled to a near-infrared charge-coupled device camera (NC70, MTI or SciCam, Scientifica). Borosilicate glass pipettes (Sutter Instruments) were pulled with resistances of 4–8MΩ. The intracellular solution for voltage-clamp recordings was (in mM): 120 CsMeSO_3_, 20 CsCl, 10 HEPES, 7 di-tris phosphocreatine, 2 MgCl_2_, 0.2 EGTA, 4 Na_2_ATP, 0.3 Tris-GTP, and in some experiments, 5 N-ethyllidocaine chloride (QX-314) (pH 7.2-7.26, 280-290 mOsm). AMPAR-mediated and NMDAR-mediated synaptic responses were evoked using a bipolar platinum/iridium electrode (FHC, CE2C275) placed in the Schaffer collaterals, ∼200 µm from the recorded cell. For synaptic experiments, a pair of 0.1 ms bipolar current pulses (50 ms ISI) was delivered via a stimulus isolation unit (Isoflex, AMPI), and stimulus intensity was adjusted to evoke a response 65-75% of the maximal response at 200 µA. After 10-20 min of stable baseline recording in the presence of picrotoxin (100 µM, PTX), netrin-1 (200 ng/ml) was bath applied for 5 min, and synaptic responses monitored for 40 min post-application. AMPAR- and NMDAR-mediated currents were evoked at −70 mV and +40 mV, respectively, during baseline conditions and 20 min post-bath application of netrin-1. Input-output tests were conducted using increasing stimulus intensity from 0-200 µA in 25 µA increments. Access and input resistances were continually monitored throughout the recording through a 50 ms, 5 mV voltage step 150 ms prior to synaptic stimulation, and data were discarded if series resistance changed >20%.

To assess HFS LTP in both NTN1 cKO and control littermates (Figure 2), as well as in the netrin-1 occlusion experiments (Figure 5), the recording mode of the cell was changed from voltage-clamp to current-clamp after a 5-10 min baseline period. Potentiation was induced by a single episode of a 1-s 100 Hz stimulation in current-clamp mode, which can elicit potentiation of evoked EPSCs at ∼150-200% of baseline values. Following stimulation, the cell was returned to voltage-clamp recording mode and held at −70 mV to record evoked responses.

For rectification experiments following bath application of exogenous netrin-1 (200 ng/ml), AMPAR-mediated currents were evoked in the presence of PTX (100 µM) and D-(-)-2-Amino-5-phosphonopentanoic acid (50 µM, D-APV) at a range of voltages (−110 mV to +50 mV in 20 mV intervals). APV was validated by measuring the lack of NMDAR-mediated currents at +40 mV. QX-314 (5 mM) was also included in the intracellular solution to prevent action potential generation. All responses were normalized to peak AMPAR-mediated current. Rectification index was computed as a ratio of current amplitude of responses evoked at −70 mV and +50 mV.

Miniature excitatory postsynaptic currents (mEPSCs) were recorded in voltage-clamp mode at a holding potential of −70 mV in the presence of PTX (100 µM) to block GABA_A_-mediated synaptic currents and tetrodotoxin (1 µM, TTX) to block sodium currents. Currents were analyzed using MiniAnalysis (Synaptosoft), and events were detected using a threshold of 7 pA (>3 pA root mean square of baseline noise levels). Cumulative distribution plots were generated using an equal number of events per condition (70 per condition), which were randomly selected from each cell, rank-ordered, and averaged across each condition.

An Axopatch 200B or 700B amplifier (Molecular Devices) were used for recordings, and signals were digitized (Digidata 1322A or 1550A, Molecular Devices). Voltage-clamp recordings were sampled at 10 kHz, filtered at 2 kHz, and acquired using pClamp software (v9.0 or v10.4, Molecular Devices).

### Hippocampal culture electrophysiology

Whole cell patch clamp recordings were made from embryonic rat hippocampal neurons (14 DIV) prepared as described above. Neurons were plated on glass coverslips at high density, and individual coverslips transferred to an upright SliceScope 2000 (Scientifica), and perfused with ACSF containing (in mM): 135 NaCl, 3.5 KCl, 1.2 MgCl_2_, 2 CaCl_2_, 10 HEPES, and 20 D-Glucose (pH: 7.4, 300 mOsm). Current clamp recordings were performed using pipettes filled with (in mM): 120 K-gluconate, 20 KCl, 10 HEPES, 7 phosphocreatine di-Tris, 2 MgCl_2_, 0.2 ethylene-glycol-bis(β-aminoethyl ether)-N,N,N’,N’-tetraacetic acid (EGTA), 4 Na-ATP, 0.3 Na-GTP (pH: 7.2-7.26, 280-290 mOsm). Access resistance was monitored throughout the recording, and cells were held at resting membrane potential. Current-clamp recordings were sampled at 20 kHz and filtered at 10 kHz using pClamp (v10.4, Molecular Devices).

### Pharmacology

Stocks were prepared at 1000x by dissolving tetrodotoxin citrate (T-550, Alomone labs), D-APV (0106, Tocris), BAPTA (ab120449, Abcam), IEM-1460 (ab141507, Abcam), Pep1-TGL (1601, Tocris), and clozapine-N-oxide (ab141704, Abcam) in water, and picrotoxin (P1675, Sigma-Aldrich) and KN62 (ab120421, Abcam) in DMSO. All other salts and drugs were from Sigma-Alrich unless otherwise noted. For certain experiments, pharmacological agents were included in the intracellular patch solution (Figures 6g-j and 7j-l). In these cases, cells were dialyzed with pipette solution for at least 15 min before beginning baseline recordings.

### Statistical analysis

Statistical analyses on parametric data were assessed using repeated measures ANOVAs, pairwise comparisons (Tukey), within-samples paired t-tests, and independent t-tests where appropriate. Normality, homoscedasticity, and outlier tests were performed on all datasets. Non-Gaussian distributed data sets were tested by Wilcoxon test for paired observations or Friedman’s repeated measures ANOVA on ranks. For mEPSC cumulative distribution data, comparisons were performed using the Kolmogorov-Smirnov test. Significant interactions and main effects were assessed using pairwise comparisons and compensated using the Tukey method for parametric data. Statistical significance was determined with *p* ≤ 0.05 using two-tailed tests. All data are presented as mean ± SEM. Data were analyzed using Clampfit 10.3 (Axon Instruments), Matlab (Mathworks), Fiji ^78^, Photoshop (Adobe), MiniAnalysis (Synaptosoft), Prism 7 (Graphpad), and Sigmaplot 11 (Systat). Plotted data were then formatted in Adobe Illustrator CS6 (Adobe Systems).

## Supplementary information

**Supplemental figure 1.**
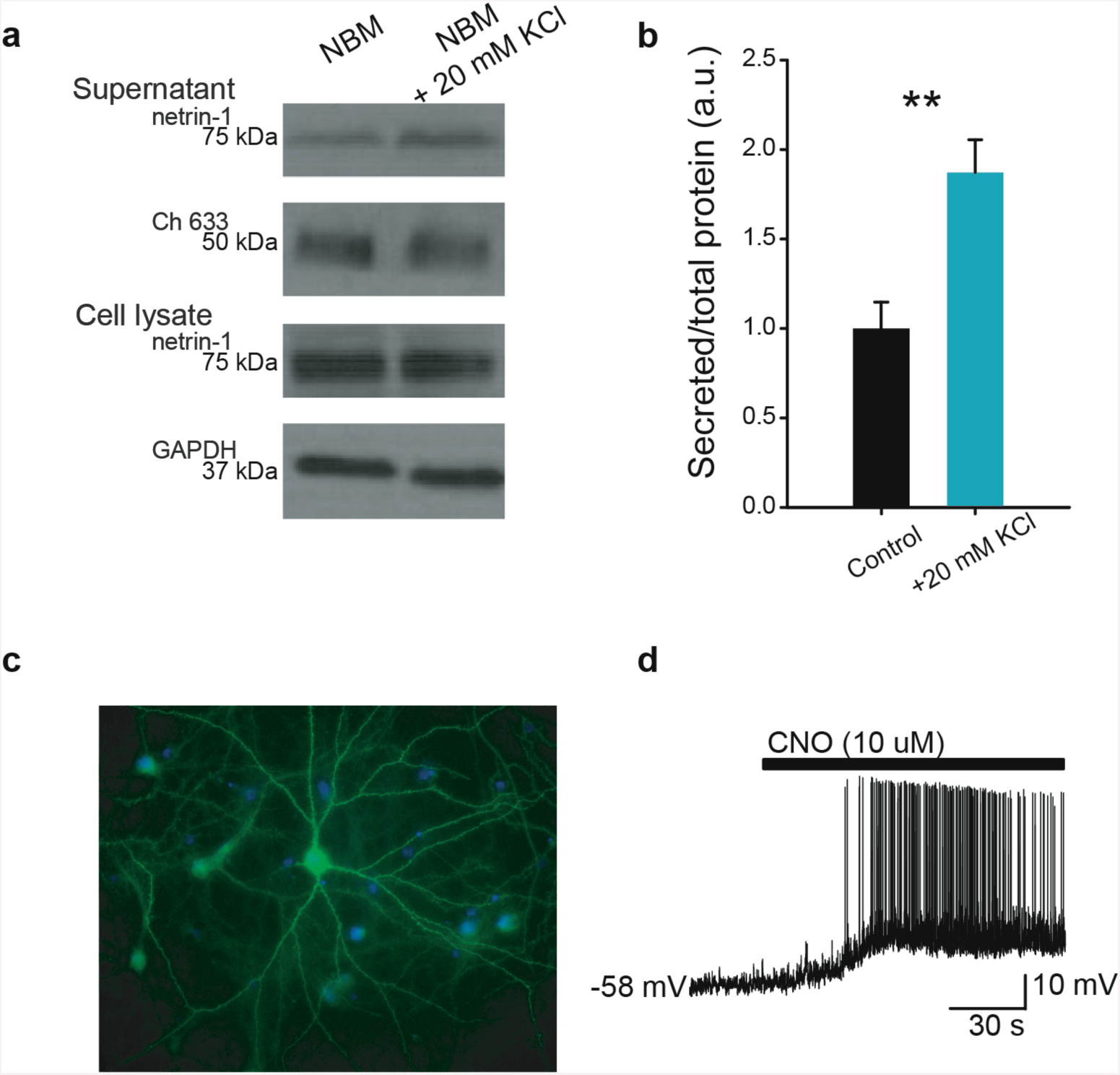
Depolarization with high KCl increases secretion of endogenous netrin-1. (**a**) Western blot showing netrin-1 protein detected in the extracellular media from dissociated hippocampal cell cultures (14 DIV) after bath application of KCl (20 mM, 20 min) or control media. Note that an Alexa 633-conjugated secondary antibody (40 ng per tube) was added to the medium as a loading control, and GAPDH was probed in cell lysate to ensure comparable cell numbers. (**b**) Group data show that 20 min bath application of media containing 20 mM KCl significantly elevates levels of netrin-1 compared to control cultures (control supernatant: 1.00±0.14, KCl supernatant: 1.87±0.18, *t*_6_ = 3.69, *p* = 0.01, independent samples *t*-test). (**c**) Photomicrograph showing a hippocampal pyramidal neuron expressing HA-tagged hM3D(Gq)-IRES-mCitrine and labelled with anti-HA (green) and Hoescht stain (blue). (**d**) Representative voltage trace showing membrane potential depolarization in a hM3D(Gq)-expressing hippocampal neuron.

**Supplemental figure 2.**
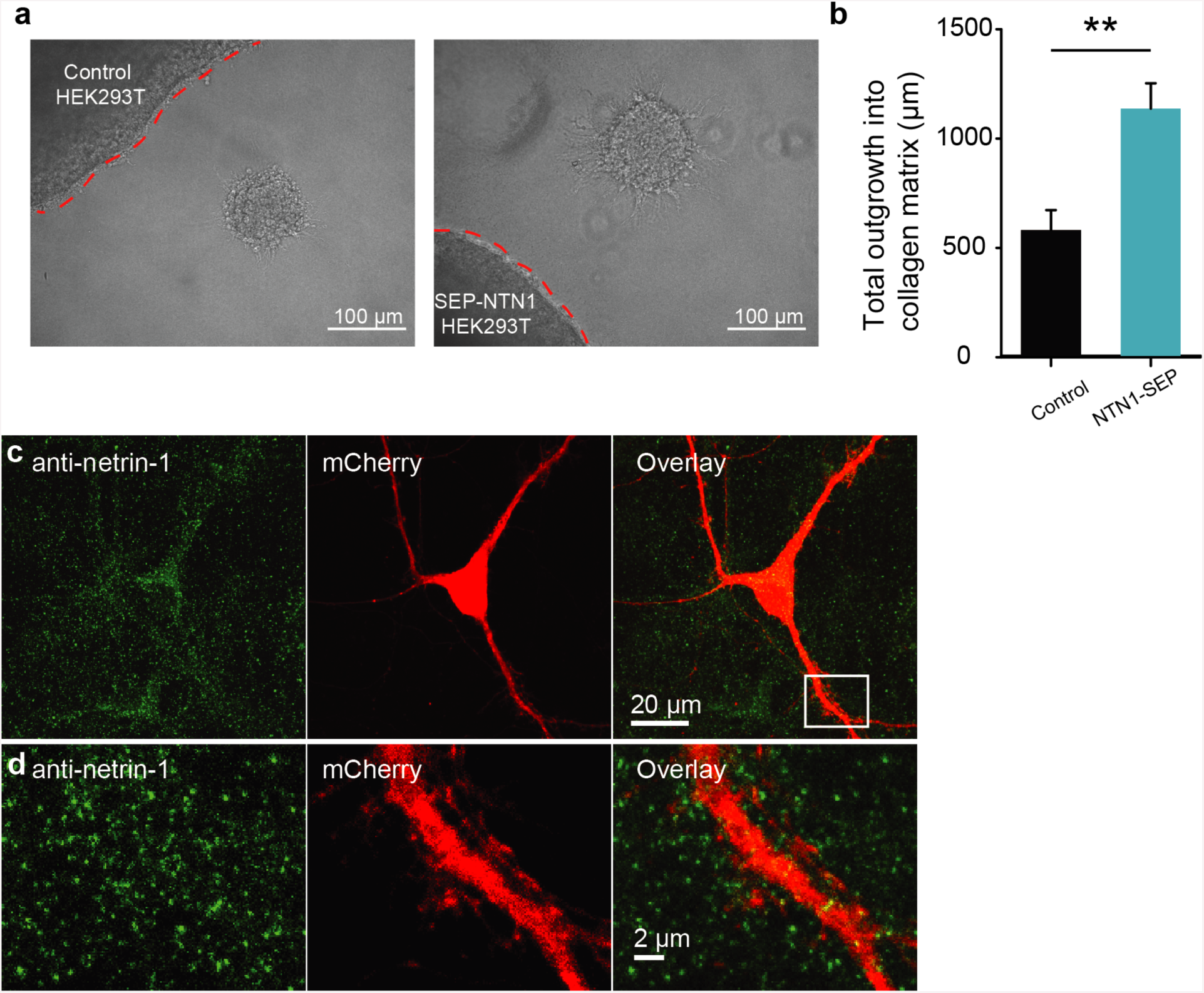
NTN1-SEP induces axon outgrowth, and shows similar distribution to endogenous netrin-1 protein in cultured hippocampal neurons. (**a**) Photomicrographs of embryonic rat dorsal spinal cord explants adjacent to control HEK293T cells (left) or expressing NTN1-SEP.**(B)**Group data show significantly higher levels of commissural axon outgrowth from explants of dorsal spinal cord when co-cultured with HEK293T cells expressing NTN1-SEP (control: 580±92, NTN1-SEP: 1137±114; *t*_10_=3.78, *p*<0.01, *n*=6 explants per condition). (**c**-**d**) Low (**c**) and high (**d**) magnification of an mCherry-expressing cultured hippocampal neuron (red) showing similar patterns of endogenous netrin-1 (green) distribution to exogenous expression of NTN1-SEP in Figure 1.

**Supplemental figure 3.**
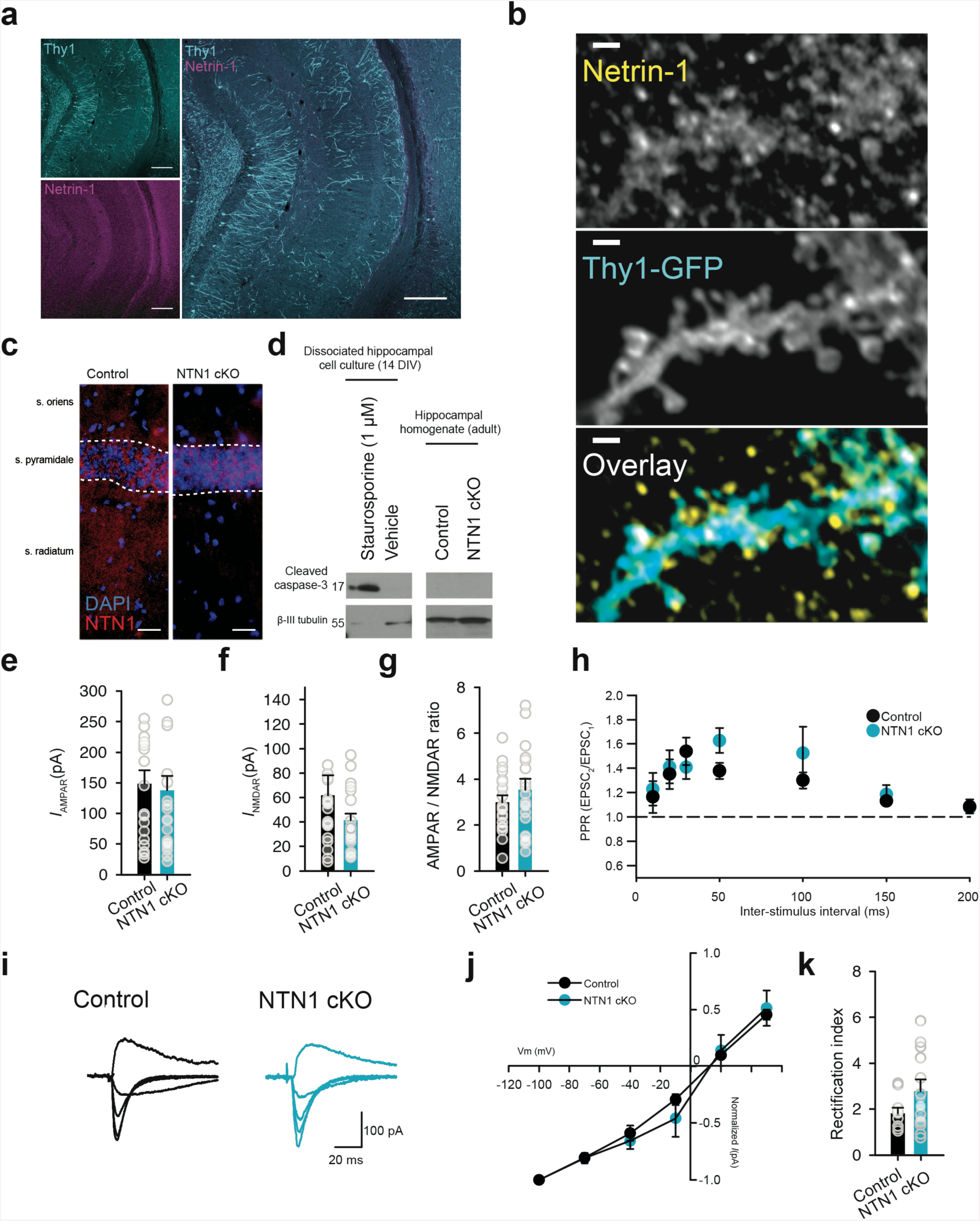
Netrin-1 is expressed in the adult hippocampus, and genetic deletion of NTN1 does not affect basal synaptic properties. (**a**) Low-(left) and high-magnification (right) images of a coronal brain section from a Thy1-GFP L15 mouse showing netrin-1 protein (red) distributed throughout the CA1 region of the hippocampus (DAPI, blue). Note netrin-1 immuno-reactivity at dendritic spines (right). Scale bar for low magnification= 200 µm. High-magnification (right) of area outlined on left (yellow square). Scale bar = 100 µm. (**b**) Representative images and overlay (right) of an apical dendrite from a Thy1-GFP-expressing CA1 pyramidal neuron costained for immunoreactivity to endogenous netrin-1 protein (yellow). Scale bar = 1 µm (**c**) Representative images of CA1 region of the hippocampus in control littermate (left) and NTN1 cKO (right, CaMKIIα∷Cre/NTN1^*fl/fl*^) showing netrin-1 immunoreactivity (red) (DAPI, blue). Scale bar = 50 µm. (**d**) Western blots of cleaved caspase-3 from dissociated hippocampal neurons (14 DIV) treated with staurosporine (1 µM, 12h; left) show increased cleaved caspase-3 compared to sister cultures treated with vehicle (right) or hippocampal homogenates from either control or T-29-CaMKIIα∷Cre/NTN1^*fl/fl*^ mice (right panel). (**e-g**) Group data show no differences between control littermates (black) or NTN1 cKO (blue) mice in raw amplitude of the AMPAR-mediated current (**e**; control: 178±32 pA, NTN1 cKO: 136±38 pA; Mann-Whitney *U* statistic=41, *p*=0.705), NMDAR-mediated current (**f**; control: 61±16 pA, NTN1 cKO: 41±5 pA; Mann-Whitney *U* statistic=162, *p*=0.316), or AMPAR-to-NMDAR ratio (**g**; control: 3.0±0.3, NTN1 cKO: 3.5±0.5; *t*_38_=0.95, *p*=0.345). Control: *n*=19 cells from 8 mice; NTN1 cKO: *n*=21 cells from 10 mice. (**h**) Effect of netrin-1 on paired-pulse ratio across a range of interstimulus intervals (10 ms: 1.52±0.19 in baseline vs. 1.30±0.12 in netrin-1; 20 ms: 1.76±0.19 in baseline vs. 1.62±0.13; 30 ms: 1.79±0.16 in baseline vs. 1.71±0.09 in netrin-1; 50 ms: 1.73±0.12 in baseline vs. 1.55±0.11 in netrin-1; 100 ms: 1.40±0.08 in baseline vs. 1.29±0.08 in netrin-1; 150 ms: 1.16±0.18 in baseline vs. 1.28±0.10 in netrin-1; 200 ms: 1.32±0.10 in baseline vs. 1.19±0.12 in netrin-1; Two-way RM-ANOVA, Main effect of ISI: *F*_6,120_=7.34, *p*<0.001, Interaction of genotype X ISI: *F*_6,120_=0.79, *p* = 0.573). (**i**) Representative traces of synaptic rectification experiments in control littermate (black) and NTN1 cKO (blue). (**j-k**) Group data show I-V curve of AMPAR-mediated synaptic responses (**j**) and rectification index (**k**; 1.82±0.24 in control littermate vs. 2.77±0.51 in NTN1 cKO, Mann-Whitney *U* statistic=45, *p*=0.339).

**Supplemental figure 4.**
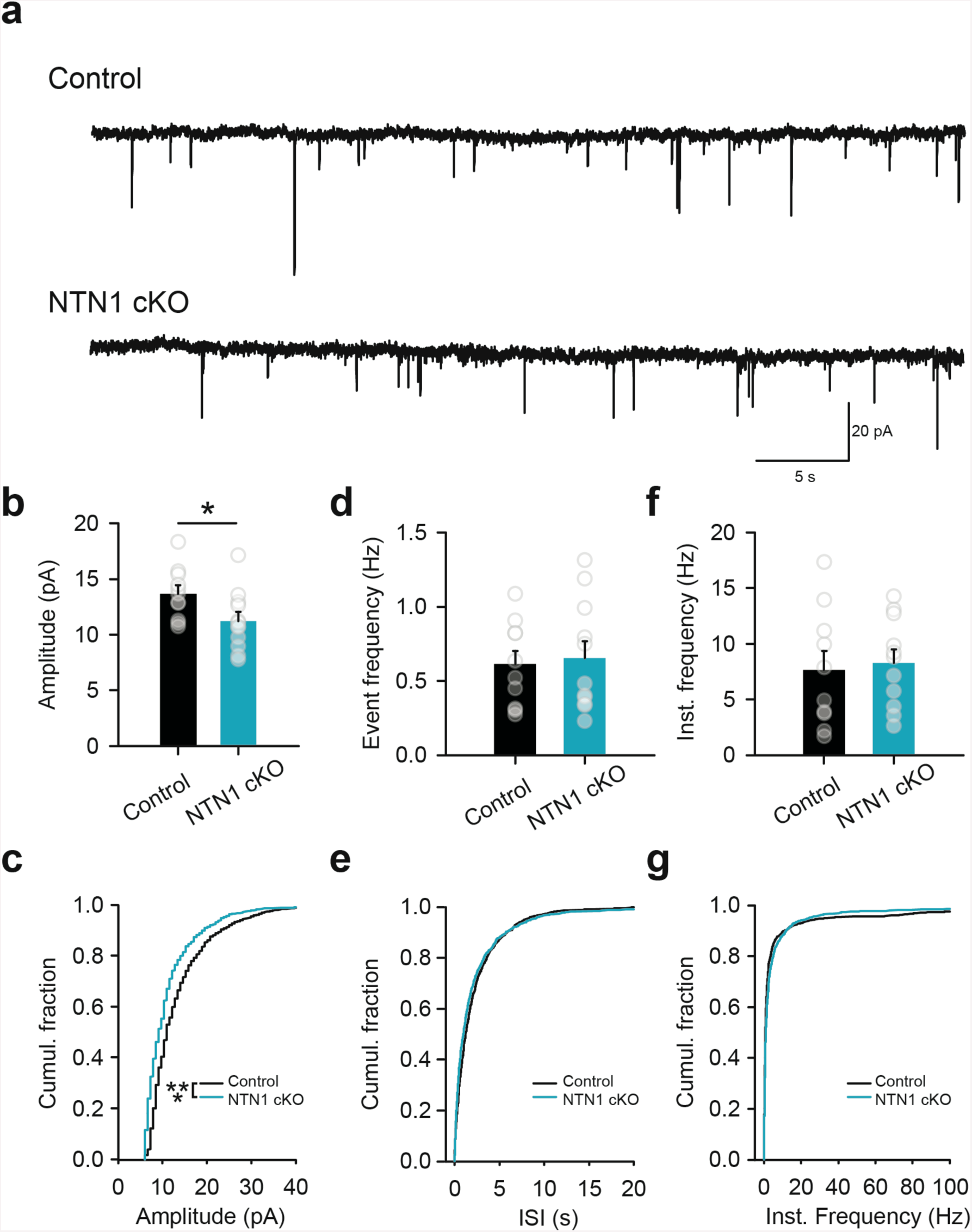
Conditional deletion of netrin-1 from forebrain excitatory neurons results in decreased miniature excitatory postsynaptic currents. **(a)** Representative traces of mEPSCs in CA1 pyramidal neurons from control littermate (top) and NTN1 cKO (bottom) mice in the presence of tetrodotoxin (1 µM) and picrotoxin (100 µM). (**b-c**) Group data (**b**) and cumulative distribution plots (**c**) show that amplitude of mEPSCs recorded from CA1 neurons of NTN1 cKO mice (blue) are significantly decreased compared to control littermates (13.6±0.7 pA in control litter-mates vs. 11.2±0.8 pA in NTN1 cKO, *t*_19_=2.18, *p*=0.042; Kolmogorov-Smirnov test for cumulative distribution data, ***: *p*<0.001). (**d**-**g**) Averaged group data (**d**, **f**) and cumulative distribution plots (**e**, **g**) show no differences between control littermates and NTN1 cKO in event (0.61±0.09 Hz in control littermates vs. 0.66±0.11 Hz in NTN1 cKO, *t*_19_=0.30, *p*=0.77) and instantaneous frequencies (7.66±1.68 Hz in control littermates vs. 8.29±1.20 Hz in NTN1 cKO, *t*_19_=0.31, *p*=0.759).

**Supplemental figure 5.**
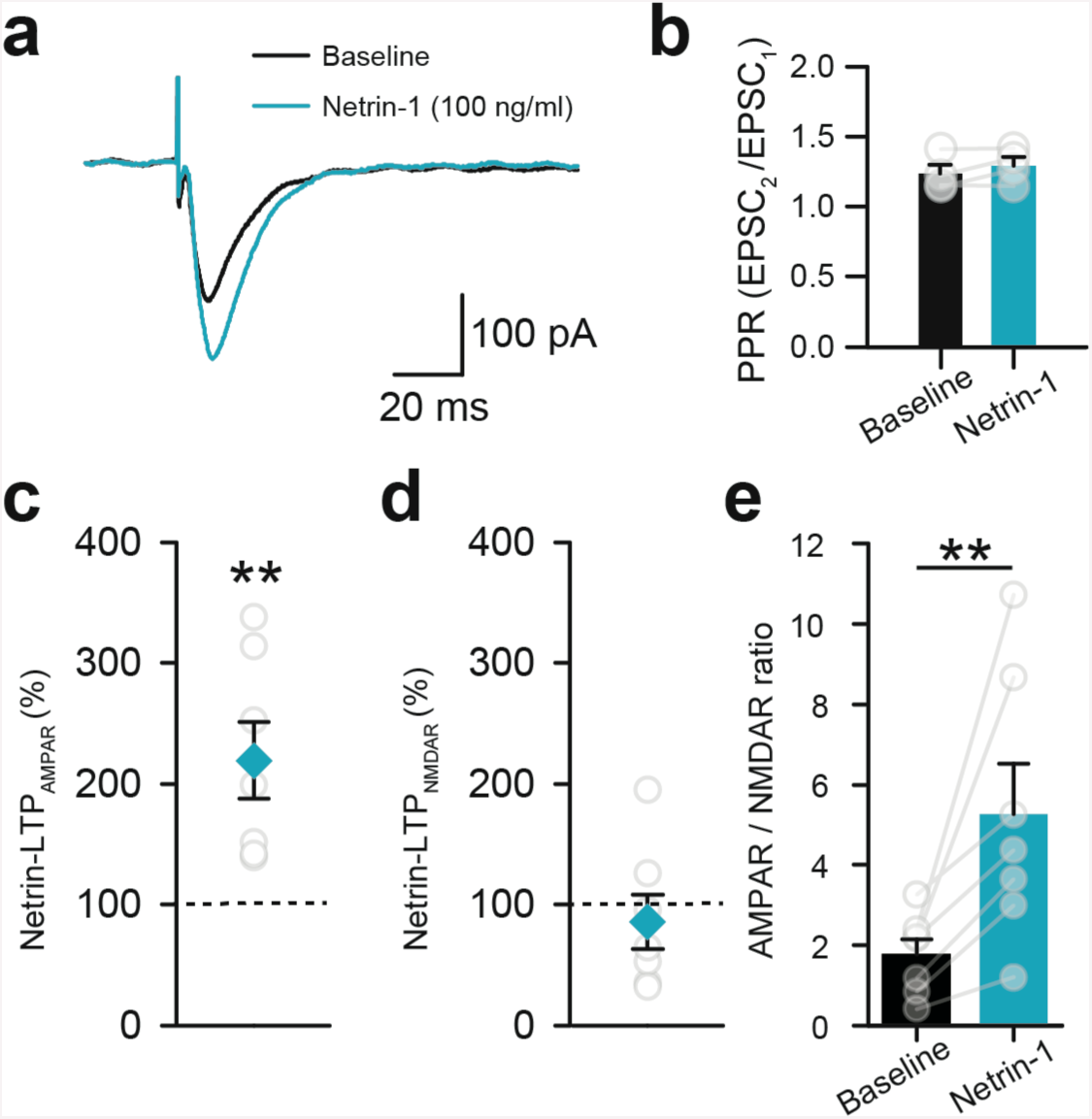
Submaximal concentrations of netrin-1 (100 ng/ml) potentiate Schaffer collateral-evoked synaptic responses in CA1 pyramidal neurons. (**a**) Representative traces of AMPAR-mediated evoked synaptic current in a CA1 pyramidal neuron in response to Schaffer collateral stimulation before (black trace) and after (blue trace) 5 min bath application of netrin-1 (100 ng/ml) in the presence of PTX (100 µM) and voltage clamped at −70 mV. (**b**) Group data show that there were no significant differences in paired-pulse ratio (50 ms ISI, *n*=4) following bath application of netrin-1 (100 ng/ml; baseline: 1.23±0.06, netrin-1: 1.29±0.06; *t*_3_=1.68, *p*=0.19, paired samples *t*-test). (**c-e**) Group data show that bath application of netrin-1 (100 ng/ml; *n*=7 cells from 5 mice) significantly enhanced AMPAR-mediated current (**c**; 219±31% of baseline, *t*_6_=4.10, *p*<0.01, paired samples *t*-test), however failed to affect NMDAR-mediated current (**d**; 85±22% of baseline, *t*_6_=1.46, *p*=0.20, paired samples *t*-test), resulting in a significant increase in the AMPAR-to-NMDAR ratio (**e**; control: 1.8±0.4, netrin-1: 5.2±1.3, *Z*=2.36, *p*<0.01, Wilcoxon Signed Rank test).

**Supplemental figure 6.**
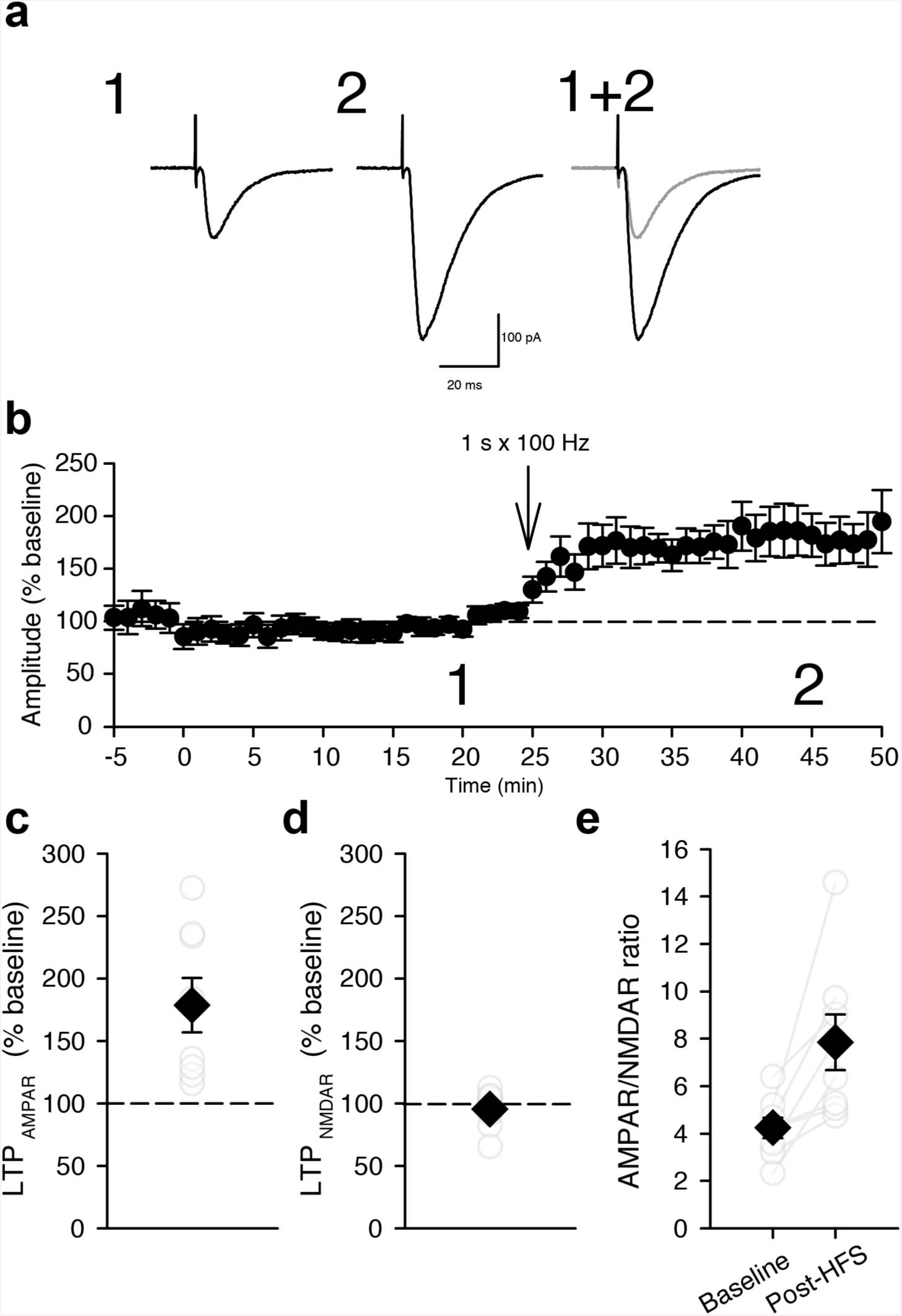
High-frequency stimulation can induce LTP after 30 min post-whole cell configuration. (**a-b**) Representative evoked AMPAR-mediated synaptic responses (**a**) and group data (**b**) during baseline (1, left in **a**) and following HFS (2, right in **a**, 1-s at 100 Hz at 30 min post-break into whole-cell configuration recording mode). (**c**-**e**) Group data show that HFS significantly increases amplitude of AMPAR-mediated currents (**c**; 178±22% of baseline, *t*_7_=3.94, *p*=0.006), but not NMDAR-mediated currents (**d**; 95±5% of baseline, *t*_7_=0.84, *p*=0.424), resulting in an increase in AMPAR-to-NMDAR ratio (**e**; 4.2±0.4 during baseline vs. 7.8±1.2 following HFS, *t*_7_=3.59, *p*=0.009).

**Supplementary movie 1.** cLTP stimulation promotes exocytosis of pH-sensitive SEP-NTN1 in hippocampal neurons. Time-lapse recording of a dendrite from a dissociated pyramidal hippocampal neuron expressing Homer1C-DsRed and SEP-NTN1 during bath perfusion of control medium (left) or cLTP-inducing solution containing 0 mM Mg^2+^, 200 µM glycine, and 10 µM bicuculline (right).

**Supplementary movie 2.** Bath application of exogenous netrin-1 (200 ng/ml) promotes synaptic insertion of GluA1. Time-lapse recording of a dissociated hippocampal pyramidal neuron (14 DIV) expressing Homer1C-DsRed (left) and SEP-GluA1 (right) before and following bath application of netrin-1 (200 ng/ml). Grey circle indicates bath application of netrin-1.

**Supplementary movie 3.** Exogenous netrin-1 promotes rapid synaptic insertion of GluA1 in hippocampal neurons. Time-lapse recording of Homer1C-DsRed (left) and SEP-GluA1 (right) at a single synapse before and following bath application of netrin-1 (200 ng/ml). Grey circle indicates bath application of netrin-1 (200 ng/ml).

## References

1. Kessels, H.W. & Malinow, R. Synaptic AMPA receptor plasticity and behavior. Neuron 61, 340–350 (2009).

2. Bosch, M. & Hayashi, Y. Structural plasticity of dendritic spines. Current opinion in neurobiology 22, 383–388 (2012).

3. Bosch, M., et al Structural and molecular remodeling of dendritic spine substructures during long-term potentiation. Neuron 82, 444–459 (2014).

4. Meyer, D., Bonhoeffer, T. & Scheuss, V. Balance and stability of synaptic structures during synaptic plasticity. Neuron 82, 430–443 (2014).

5. Bourne, J. & Harris, K.M. Do thin spines learn to be mushroom spines that remember? Current opinion in neurobiology 17, 381–386 (2007).

6. Kasai, H., Fukuda, M., Watanabe, S., Hayashi-Takagi, A. & Noguchi, J. Structural dynamics of dendritic spines in memory and cognition. Trends Neurosci 33, 121–129 (2010).

7. Goldman, J.S., et al Netrin-1 promotes excitatory synaptogenesis between cortical neurons by initiating synapse assembly. The Journal of neuroscience: the official journal of the Society for Neuroscience 33, 17278–17289 (2013).

8. Kennedy, T.E., Serafini, T., de la Torre, J.R. & Tessier-Lavigne, M. Netrins are diffusible chemotropic factors for commissural axons in the embryonic spinal cord. Cell 78, 425–435 (1994).

9. Serafini, T., et al The netrins define a family of axon outgrowth-promoting proteins homologous to C. elegans UNC-6. Cell 78, 409–424 (1994).

10. Lai Wing Sun, K., Correia, J.P. & Kennedy, T.E. Netrins: versatile extracellular cues with diverse functions. Development 138, 2153–2169 (2011).

11. Tada, T. & Sheng, M. Molecular mechanisms of dendritic spine morphogenesis. Current opinion in neurobiology 16, 95–101 (2006).

12. Colon-Ramos, D.A., Margeta, M.A. & Shen, K. Glia promote local synaptogenesis through UNC-6 (netrin) signaling in C. elegans. Science 318, 103–106 (2007).

13. Poon, V.Y., Klassen, M.P. & Shen, K. UNC-6/netrin and its receptor UNC-5 locally exclude presynaptic components from dendrites. Nature 455, 669–673 (2008).

14. Kim, S. & Martin, K.C. Neuron-wide RNA transport combines with netrin-mediated local translation to spatially regulate the synaptic proteome. eLife 4 (2015).

15. Horn, K.E., et al DCC expression by neurons regulates synaptic plasticity in the adult brain. Cell reports 3, 173–185 (2013).

16. Ahmed, G., et al Draxin inhibits axonal outgrowth through the netrin receptor DCC. The Journal of neuroscience: the official journal of the Society for Neuroscience 31, 14018–14023 (2011).

17. Haddick, P.C., et al Defining the ligand specificity of the deleted in colorectal cancer (DCC) receptor. PLoS One 9, e84823 (2014).

18. Wei, P., et al The Cbln family of proteins interact with multiple signaling pathways. J Neurochem 121, 717–729 (2012).

19. Ryan, T.A. & Smith, S.J. Vesicle Pool Mobilization during Action-Potential Firing at Hippocampal Synapses. Neuron 14, 983–989 (1995).

20. Yudowski, G.A., et al Real-time imaging of discrete exocytic events mediating surface delivery of AMPA receptors. The Journal of neuroscience: the official journal of the Society for Neuroscience 27, 11112–11121 (2007).

21. Bliss, T.V. & Collingridge, G.L. Expression of NMDA receptor-dependent LTP in the hippocampus: bridging the divide. Molecular brain 6, 5 (2013).

22. Lu, W., et al Activation of synaptic NMDA receptors induces membrane insertion of new AMPA receptors and LTP in cultured hippocampal neurons. Neuron 29, 243–254 (2001).

23. Lemieux, M., et al Translocation of CaMKII to dendritic microtubules supports the plasticity of local synapses. J Cell Biol 198, 1055–1073 (2012).

24. Cembrowski, M.S., Wang, L., Sugino, K., Shields, B.C. & Spruston, N. Hipposeq: a comprehensive RNA-seq database of gene expression in hippocampal principal neurons. eLife 5 (2016).

25. Bin, J.M., et al Complete Loss of Netrin-1 Results in Embryonic Lethality and Severe Axon Guidance Defects without Increased Neural Cell Death. Cell reports 12, 1099–1106 (2015).

26. Malenka, R.C. & Bear, M.F. LTP and LTD: an embarrassment of riches. Neuron 44, 5–21 (2004).

27. Tsien, J.Z., et al Subregion- and cell type-restricted gene knockout in mouse brain. Cell 87, 1317–1326 (1996).

28. Forcet, C., et al The dependence receptor DCC (deleted in colorectal cancer) defines an alternative mechanism for caspase activation. Proc Natl Acad Sci U S A 98, 3416–3421 (2001).

29. Stavoe, A.K. & Colon-Ramos, D.A. Netrin instructs synaptic vesicle clustering through Rac GTPase, MIG-10, and the actin cytoskeleton. J Cell Biol 197, 75–88 (2012).

30. Lisman, J. & Raghavachari, S. A unified model of the presynaptic and postsynaptic changes during LTP at CA1 synapses. Science’s STKE: signal transduction knowledge environment 2006, re11 (2006).

31. Morita, D., Rah, J.C. & Isaac, J.T. Incorporation of inwardly rectifying AMPA receptors at silent synapses during hippocampal long-term potentiation. Philosophical transactions of the Royal Society of London. Series B, Biological sciences 369, 20130156 (2014).

32. Keino-Masu, K., et al Deleted in Colorectal Cancer (DCC) encodes a netrin receptor. Cell 87, 175–185 (1996).

33. Opazo, P., et al CaMKII triggers the diffusional trapping of surface AMPARs through phosphorylation of stargazin. Neuron 67, 239–252 (2010).

34. Hayashi, Y., et al Driving AMPA receptors into synapses by LTP and CaMKII: requirement for GluR1 and PDZ domain interaction. Science 287, 2262–2267 (2000).

35. Wang, G.X. & Poo, M.M. Requirement of TRPC channels in netrin-1-induced chemotropic turning of nerve growth cones. Nature 434, 898–904 (2005).

36. Hollmann, M., Hartley, M. & Heinemann, S. Ca2+ permeability of KA-AMPA--gated glutamate receptor channels depends on subunit composition. Science 252, 851–853 (1991).

37. Burnashev, N., Monyer, H., Seeburg, P.H. & Sakmann, B. Divalent ion permeability of AMPA receptor channels is dominated by the edited form of a single subunit. Neuron 8, 189–198 (1992).

38. Granger, A.J., Shi, Y., Lu, W., Cerpas, M. & Nicoll, R.A. LTP requires a reserve pool of glutamate receptors independent of subunit type. Nature 493, 495–500 (2013).

39. Jonas, P. & Burnashev, N. Molecular mechanisms controlling calcium entry through AMPA-type glutamate receptor channels. Neuron 15, 987–990 (1995).

40. Luscher, C. & Malenka, R.C. NMDA receptor-dependent long-term potentiation and long-term depression (LTP/LTD). Cold Spring Harb Perspect Biol 4 (2012).

41. Edelmann, E., et al Theta Burst Firing Recruits BDNF Release and Signaling in Postsynaptic CA1 Neurons in Spike-Timing-Dependent LTP. Neuron 86, 1041–1054 (2015).

42. Herron, C.E., Lester, R.A., Coan, E.J. & Collingridge, G.L. Frequency-dependent involvement of NMDA receptors in the hippocampus: a novel synaptic mechanism. Nature 322, 265–268 (1986).

43. Regehr, W.G. Short-term presynaptic plasticity. Cold Spring Harb Perspect Biol 4, a005702 (2012).

44. Gu, Y., et al Differential vesicular sorting of AMPA and GABAA receptors. Proc Natl Acad Sci U S A 113, E922–931 (2016).

45. Lledo, P.M., Zhang, X., Sudhof, T.C., Malenka, R.C. & Nicoll, R.A. Postsynaptic membrane fusion and long-term potentiation. Science 279, 399–403 (1998).

46. Xiao, B., et al Homer regulates the association of group 1 metabotropic glutamate receptors with multivalent complexes of homer-related, synaptic proteins. Neuron 21, 707–716 (1998).

47. Brakeman, P.R., et al Homer: a protein that selectively binds metabotropic glutamate receptors. Nature 386, 284–288 (1997).

48. Berry, K.P. & Nedivi, E. Spine Dynamics: Are They All the Same? Neuron 96, 43–55 (2017).

49. Zito, K., Scheuss, V., Knott, G., Hill, T. & Svoboda, K. Rapid functional maturation of nascent dendritic spines. Neuron 61, 247–258 (2009).

50. Lambert, J.T., Hill, T.C., Park, D.K., Culp, J.H. & Zito, K. Protracted and asynchronous accumulation of PSD95-family MAGUKs during maturation of nascent dendritic spines. Dev Neurobiol 77, 1161–1174 (2017).

51. El-Husseini, A.E., Schnell, E., Chetkovich, D.M., Nicoll, R.A. & Bredt, D.S. PSD-95 involvement in maturation of excitatory synapses. Science 290, 1364–1368 (2000).

52. Zhou, Z., et al The C-terminal tails of endogenous GluA1 and GluA2 differentially contribute to hippocampal synaptic plasticity and learning. Nat Neurosci 21, 50–62 (2018).

53. Benke, T.A., Luthi, A., Isaac, J.T. & Collingridge, G.L. Modulation of AMPA receptor unitary conductance by synaptic activity. Nature 393, 793–797 (1998).

54. Manabe, T., Renner, P. & Nicoll, R.A. Postsynaptic contribution to long-term potentiation revealed by the analysis of miniature synaptic currents. Nature 355, 50–55 (1992).

55. Oliet, S.H., Malenka, R.C. & Nicoll, R.A. Bidirectional control of quantal size by synaptic activity in the hippocampus. Science 271, 1294–1297 (1996).

56. Bourne, J.N. & Harris, K.M. Coordination of size and number of excitatory and inhibitory synapses results in a balanced structural plasticity along mature hippocampal CA1 dendrites during LTP. Hippocampus 21, 354–373 (2011).

57. Kerchner, G.A. & Nicoll, R.A. Silent synapses and the emergence of a postsynaptic mechanism for LTP. Nat Rev Neurosci 9, 813–825 (2008).

58. Plant, K., et al Transient incorporation of native GluR2-lacking AMPA receptors during hippocampal long-term potentiation. Nat Neurosci 9, 602–604 (2006).

59. Sutton, M.A., et al Miniature neurotransmission stabilizes synaptic function via tonic suppression of local dendritic protein synthesis. Cell 125, 785–799 (2006).

60. Herring, B.E. & Nicoll, R.A. Long-Term Potentiation: From CaMKII to AMPA Receptor Trafficking. Annu Rev Physiol 78, 351–365 (2016).

61. Lee, S.J., Escobedo-Lozoya, Y., Szatmari, E.M. & Yasuda, R. Activation of CaMKII in single dendritic spines during long-term potentiation. Nature 458, 299–304 (2009).

62. Matsuzaki, M., et al Dendritic spine geometry is critical for AMPA receptor expression in hippocampal CA1 pyramidal neurons. Nat Neurosci 4, 1086–1092 (2001).

63. Beique, J.C., et al Synapse-specific regulation of AMPA receptor function by PSD-95. Proc Natl Acad Sci U S A 103, 19535–19540 (2006).

64. Murakoshi, H., Wang, H. & Yasuda, R. Local, persistent activation of Rho GTPases during plasticity of single dendritic spines. Nature 472, 100–104 (2011).

65. Matsuzaki, M., Honkura, N., Ellis-Davies, G.C. & Kasai, H. Structural basis of long-term potentiation in single dendritic spines. Nature 429, 761–766 (2004).

66. Shekarabi, M., et al Deleted in colorectal cancer binding netrin-1 mediates cell substrate adhesion and recruits Cdc42, Rac1, Pak1, and N-WASP into an intracellular signaling complex that promotes growth cone expansion. The Journal of neuroscience: the official journal of the Society for Neuroscience 25, 3132–3141 (2005).

67. DeGeer, J., et al Tyrosine phosphorylation of the Rho guanine nucleotide exchange factor Trio regulates netrin-1/DCC-mediated cortical axon outgrowth. Mol Cell Biol 33, 739–751 (2013).

68. Moore, S.W., et al Rho inhibition recruits DCC to the neuronal plasma membrane and enhances axon chemoattraction to netrin 1. Development 135, 2855–2864 (2008).

69. Meng, Y., Zhang, Y. & Jia, Z. Synaptic transmission and plasticity in the absence of AMPA glutamate receptor GluR2 and GluR3. Neuron 39, 163–176 (2003).

70. Park, P., et al Calcium-Permeable AMPA Receptors Mediate the Induction of the Protein Kinase A-Dependent Component of Long-Term Potentiation in the Hippocampus. The Journal of neuroscience: the official journal of the Society for Neuroscience 36, 622–631 (2016).

71. Hong, K., Nishiyama, M., Henley, J., Tessier-Lavigne, M. & Poo, M. Calcium signalling in the guidance of nerve growth by netrin-1. Nature 403, 93–98 (2000).

72. Nishiyama, M., Hong, K., Mikoshiba, K., Poo, M.M. & Kato, K. Calcium stores regulate the polarity and input specificity of synaptic modification. Nature 408, 584–588 (2000).

73. Blasiak, A., Kilinc, D. & Lee, G.U. Neuronal Cell Bodies Remotely Regulate Axonal Growth Response to Localized Netrin-1 Treatment via Second Messenger and DCC Dynamics. Front Cell Neurosci 10, 298 (2016).

74. De Paola, V., Arber, S. & Caroni, P. AMPA receptors regulate dynamic equilibrium of presynaptic terminals in mature hippocampal networks. Nat Neurosci 6, 491–500 (2003).

75. Lavoie-Cardinal, F., Salesse, C., Bergeron, E., Meunier, M. & De Koninck, P. Gold nanoparticle-assisted all optical localized stimulation and monitoring of Ca(2)(+) signaling in neurons. Scientific reports 6, 20619 (2016).

76. Bouchard, J.F., Horn, K.E., Stroh, T. & Kennedy, T.E. Depolarization recruits DCC to the plasma membrane of embryonic cortical neurons and enhances axon extension in response to netrin-1. J Neurochem 107, 398–417 (2008).

77. Gabriel, L.R., Wu, S. & Melikian, H.E. Brain slice biotinylation: an ex vivo approach to measure region-specific plasma membrane protein trafficking in adult neurons. Journal of visualized experiments: JoVE (2014).

78. Schindelin, J., et al Fiji: an open-source platform for biological-image analysis. Nat Methods 9, 676–682 (2012).

79. Ng, M., et al Transmission of olfactory information between three populations of neurons in the antennal lobe of the fly. Neuron 36, 463–474 (2002).

80. Hudmon, A., et al A mechanism for Ca2+/calmodulin-dependent protein kinase II clustering at synaptic and nonsynaptic sites based on self-association. The Journal of neuroscience: the official journal of the Society for Neuroscience 25, 6971–6983 (2005).

